# Global RNA Virome in Wastewater Treatment Plants Reveals Ecological Insights and Human Health Implications

**DOI:** 10.1101/2024.03.12.584551

**Authors:** Ling Yuan, Lin-xing Chen, Hanqing Yu, Jizhong Zhou, Ren Sun, Feng Ju

## Abstract

RNA viruses are widely recognized for their roles in causing human diseases and shaping Earth’s biodiversity. Wastewater treatment plants (WWTPs) are eco-friendly biotechnological systems where the roles of RNA viruses in process engineering and sanitation remain unclear. This study analyzed RNA sequencing dataset (> 3.8 Tb) from global WWTPs to examine the diversity, host associations, and auxiliary metabolic functions of RNA viruses. We identified 11,414 RNA virus operational taxonomic units (vOTUs), expanding the known diversity of RNA viruses in WWTPs by 67%. The RNA viral community in WWTPs was dominated by prokaryotic viruses, including both established RNA phage lineages and novel clades with broad ecological distributions, highlighting their underestimated diversity and broad niche breadths. Notably, a vOTU from the base*-Howeltoviricetes* phage clade was associated with the pathogenic bacterium *Aliarcobacter cryaerophilus*, suggesting potential applications in RNA phage therapy. Furthermore, the examined distribution and fate of human RNA viruses emphasized the utility of quantitative metatranscriptomics-based wastewater surveillance for public health monitoring. The discovery of auxiliary metabolic genes encoded by RNA viruses further revealed their involvement in critical host metabolic pathways such as translation and cellular respiration. These findings underscore the multifaceted roles of RNA viruses in the critical engineered systems.

## Introduction

RNA viruses are abundant and influential biological entities that span a wide range of ecosystems, infecting both eukaryotes and prokaryotes. These viruses play critical roles in human health^1–3^, host metabolism manipulation^4–6^, and the maintenance of global biodiversity^7–9^. Recent breakthroughs in RNA virus research, driven by the detection of RNA-dependent RNA polymerase (RdRp) in metatranscriptomic datasets, have dramatically expanded our understanding of their diversity^6–8, 10–16^. Metatranscriptomics has emerged as a powerful tool for exploring RNA viruses across diverse ecosystems ^12, 17^. Recent large-scale surveys of RNA viruses in marine^6, 10, 11^ and soil ecosystems^13, 18^ reveal that RNA viruses are integral components of biodiversity in both aquatic and terrestrial ecosystems.

Wastewater treatment plants (WWTPs) are among the world’s most significant biotechnological systems, designed to safeguard human and environmental health. Each year, WWTPs process and purify over 180 km^3^ of wastewater, leveraging the activities of microbial communities to remove organic pollutants and nutrients^19, 20^. As such, WWTPs serve as both physical and ecological bridges between human populations and natural ecosystems^21^. Both DNA and RNA viruses are fundamental components of wastewater ecosystems, influencing microbial community dynamics, biogeochemical cycles, and public health. While notable progress has been made in understanding the diversity of DNA viruses in WWTPs^22, 23^, our knowledge of RNA viruses in WWTPs remains limited.

The RNA virosphere in wastewater is highly diverse, encompassing not only prokaryotic RNA viruses but also human RNA viruses originating from feces and urine origins, as well as RNA viruses associated with animals, plants, protozoa, and fungi^24^. RNA phages are key players in WWTP ecosystems where they infect and lyse bacterial hosts, thereby regulating bacterial populations and influencing the structure of microbial communities essential for wastewater treatment. Understanding the dynamics of these RNA viruses including the identification and characterization of specific clades and novel lineages, is crucial for deciphering their role in WWTP processes. Simultaneously, human RNA viruses in wastewater have become critical tools in public health surveillance. The application of wastewater-based epidemiology (WBE), exemplified during the COVID-19 pandemic, has proven invaluable in tracking population-wide health trends and enabling early outbreak detection^25, 26^. Despite these advancements, a comprehensive understanding of the spectrum of human RNA viruses present in WWTPs is still lacking. Furthermore, while auxiliary metabolic genes (AMGs) have been identified in DNA viruses and shown to influence host metabolic pathways ^22, 27–30^, the extent to which RNA viruses contribute to host metabolism and biochemical cycling through AMGs remains unclear. Addressing these knowledge gaps on the diversity, host interactions, biochemical roles, and fate of RNA viruses in WWTPs could uncover novel ecological functions and health implications of RNA viruses.

To bridge these knowledge gaps, we conducted an extensive study to investigate RNA viruses and their ecological roles in WWTPs worldwide. By analyzing the largest metatranscriptomic dataset of WWTPs to date (>3.8 Tb), we created the WWTP RNA Viruses Database (WRVirDB). This effort identified abundant RNA phages in WWTPs, including both known and newly discovered clades. Our study also explores the presence of RNA virus-encoded AMGs, revealing their potential ecological significance. Additionally, we mapped the global distribution of human RNA viruses in wastewater and identified novel viral species, highlighting the utility of wastewater-based surveillance for environmental epidemiology and public health management.

## Results

### Metatranscriptomic profiling and cross-compartment distribution patterns of RNA viruses in the global WWTPs

To explore the diversity of RNA viruses in global WWTPs, we analyzed > 3.8 Tb of RNA sequencing data from 557 publicly available metatranscriptomes spanning five WWTP compartments (Fig. 1a), including influent (IN), denitrification sludge (DS), activated sludge (AS), effluent (EF), and anaerobic digestion sludge (AD). A total of 55,900 RNA viral contigs were identified by iteratively hidden Markov model (HMM)-based searching of RdRp genes and subsequent assessment of RNA virus authenticity (See Methods and Extended Data Fig. 1). These RNA viral contigs were further clustered into 11,414 RNA virus operational taxonomic units (vOTUs) at an average nucleotide identity (ANI) of 90% over 80% alignment fraction of the shorter sequence (AF) to empirically evaluate RNA viruses at the “species” rank (Supplementary Table 2). Of these RNA vOTUs, 9,276 encoded “complete” (or near-complete) RdRp domains. Although the recovery of RNA viral contigs was gradually to be saturated after eight iterative searches, the overall diversity of unique RNA vOTUs in the WWTPs showed no indication of reaching saturation (Fig. 1b), suggesting that additional samples are required to assemble a more comprehensive catalog of RNA viruses in WWTP systems.

**Fig. 1.**
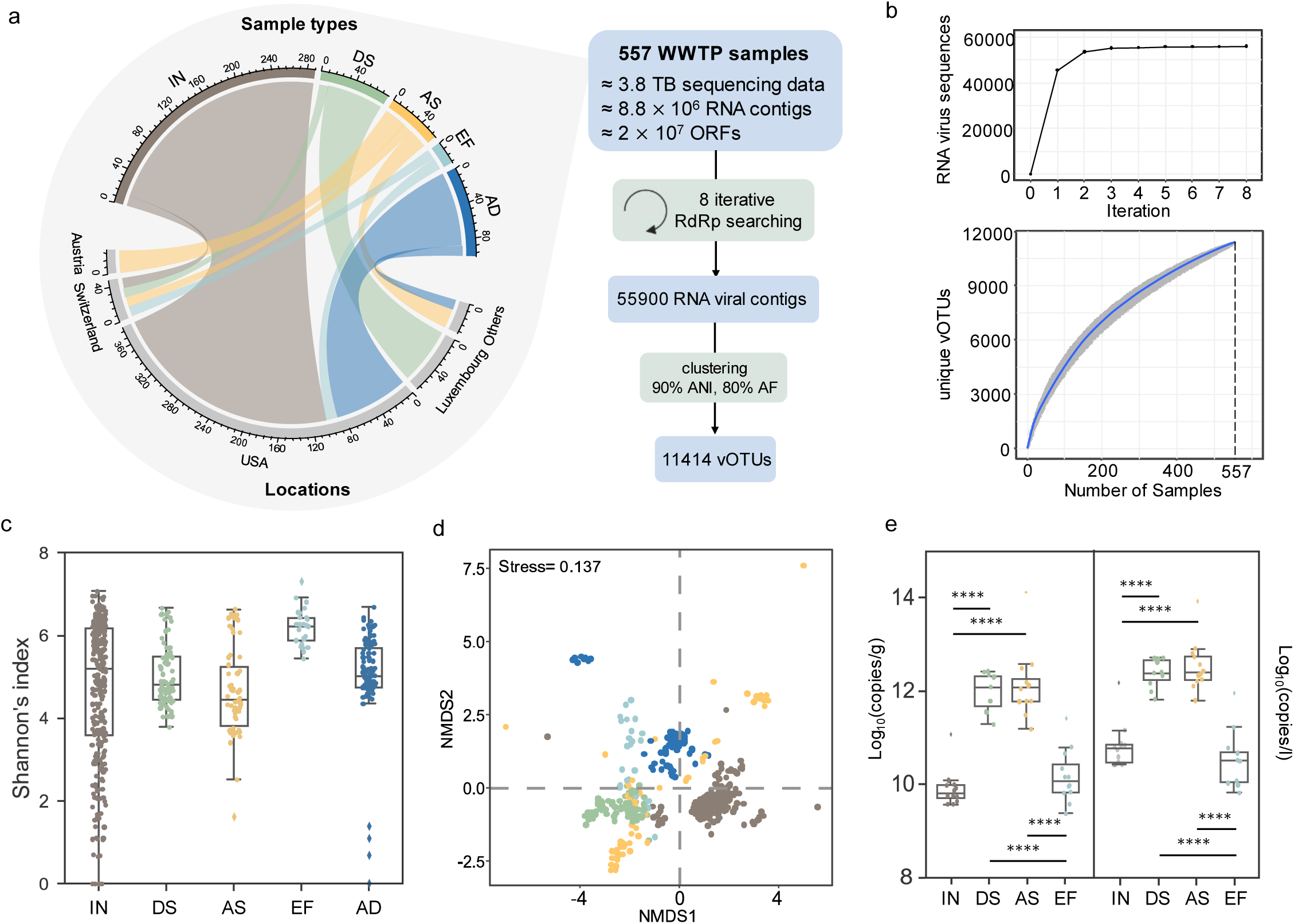
Exploring RNA viruses in the global WWTP metatranscriptomes. a. Distribution of sample types and locations of 557 publicly available WWTP metatranscriptomes and the brief bioinformatic workflow for RNA vOTUs identification. b. The number of RNA viral contigs detected across eight hmmsearch iterations (upper panel) and the accumulation curve of unique RNA vOTUs identified in the samples (below panel). c. The α-diversity of RNA vOTUs measured using Shannon’s index across five sample types in all WWTP samples. d. NMDS analysis of abundance distribution of RNA vOTUs in all WWTP samples. e. The absolute abundance of all identified RNA vOTUs expressed as copies per gram biomass (left) and copies per liter (right), is shown for four sample types in the 47 WWTP samples from Switzerland. Statistical significance was determined using the Mann-Whitney U test, with significance levels represented as ∗∗∗∗ for adjusted *P* < 0.00001. IN: influent, DS: denitrification sludge, AS: activated sludge, EF: effluent, AD: anaerobic digester sludge.

Further analysis of α-diversity patterns of RNA vOTUs revealed a highly variable richness of RNA viruses in the WWTP samples (Fig. 1c). Non-Metric Multidimensional Scaling (NMDS) analysis based on vOTUs composition demonstrated a clear separation of RNA viral communities by sample type (ANOSIM R^2^ = 0.821, *P* = 0.001; Fig. 1d). To estimate the absolute concentration of RNA vOTUs, we analyzed 47 samples from Switzerland spiked with known copies of mRNA internal standards^31^. The absolute concentration of RNA vOTUs reached ∼ 10^12^ copies/gram-biomass in sludge samples and ∼ 10^10^ copies/gram-biomass in influent and effluent wastewater samples (Fig. 1e). Notably, the effluent samples from Switzerland displayed higher variations in terms of α-diversity and combined absolute concentration of RNA vOTUs compared to the related influent samples (Supplementary Fig. 1 and Fig. 1e), indicating that wastewater treatment processes may exert uneven filtering or removal effects on the RNA virome, leading to more dispersed RNA viral community structure in the effluent (Supplementary Text 1).

### Knowledge expansion of phylogenetic diversity and taxonomic novelty of RNA viruses in the global WWTPs

RdRp-based megataxon clustering and phylogeny analysis supported the five established RNA virus phyla approved by the International Committee on Taxonomy of Viruses (ICTV) (Supplementary Table 4, Fig. 2a and Supplementary Fig. 2). Most of the RNA vOTUs that encoded “complete” RdRp domains (8,755 / 9,276) can be posited into already established phyla. Moreover, 5,945 RNA vOTUs could be classified into known viral families. The RNA vOTUs were mostly taxonomically assigned to *Fiersviridae* (1,772) and *Steitzviridae* (580), both of which were known viral families infecting prokaryotes, followed by *Botourmiaviridae* (488), which was known to infect plants or fungi (Extended Data Fig. 2).

**Fig. 2.**
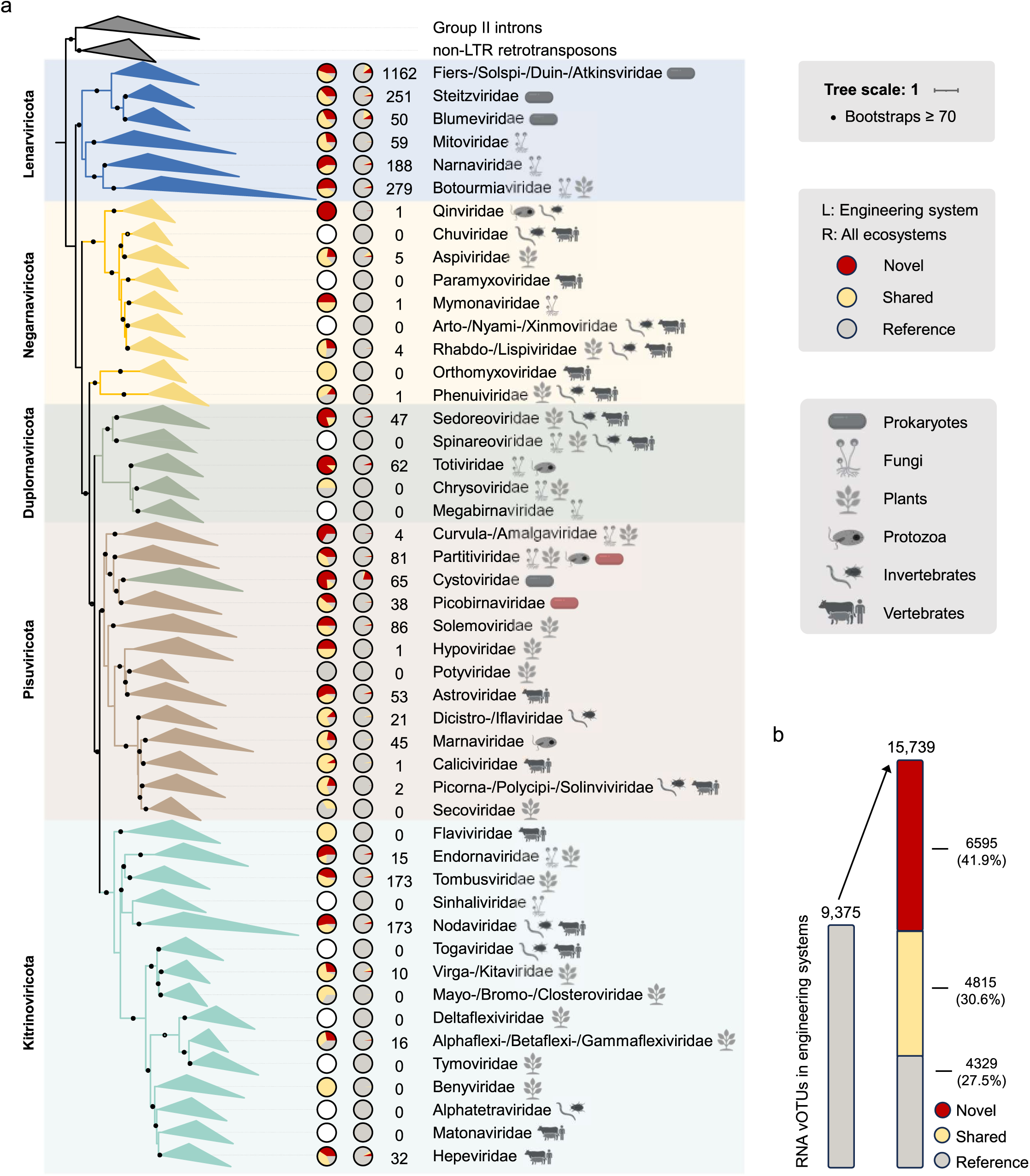
The phylogenetic and taxonomical expansion of RNA viral diversity. a. The global RdRp-based phylogenetic tree. The tree consists of reference RdRp centroids and vOTUs encoded “complete” RdRp domains, classified into known RNA viral families. Each family clade is accompanied by two pie charts, representing the distribution of its clusters within the GRvOTU set that occurred in WWTP systems (left pie) and all ecosystems (right pie). In each pie chart, the segments indicate the proportions of novel (members only identified in the WWTPs), reference (members only identified in the reference dataset), and shared (members present in both datasets) clusters. The numbers adjacent to the pie charts show the count of ‘novel’ clusters within the GRvOTU set for the family. Host range icons next to family names represent their known host ranges. Icons highlighted in red indicate a subset of *Partitiviridae* and *Picobirnaviridae* identified as bacteriophages, challenging previously assumed host ranges (see Section: Tracking Host Associations of RNA Prokaryotic Viruses in Global WWTPs). b. Expansion of RNA viral diversity in engineering systems, showing the number of known RNA viral clusters (left) and the distribution of ‘novel’, ‘shared’, and ‘reference’ clusters (right) within the GRvOTU set.

To assess our expansion of RNA virus diversity in WWTPs, the RNA viruses identified in this study were clustered with the RNA viral sequences from the NCBI Virus database and recent RNA virome studies^10–13, 18^. Clustering was conducted at both the genome level (GRvOTU set) and the RdRp gene level (RCR90 set). Our findings substantially increased the diversity of unique RNA viral clusters in WWTP systems, from 9,375 previously known RNA viral clusters to a total of 15,739 RNA viral clusters (Fig. 2b). At the viral family level, clustering statistics based on genome and RdRp gene data were highly consistent (Supplementary Table 5). Although WWTP RNA viruses showed a limited impact on expanding the overall diversity for RNA viral families, they significantly enriched the diversity of bacteriophages within WWTP systems (Fig. 2a and Supplementary Table 5). *Leviviricetes* and *Cystoviridae* are the only two groups known to infect bacteria so far. The WWTP RNA viruses discovered in this study contributed 32.4% to 73.0% of novel diversity to RNA viral families within these two clades in WWTP systems (Extended Data Fig. 3).

The RNA viruses identified in this study also supported the establishment of potential new clades. A substantial number of RNA viruses encoding “complete” RdRp could not be classified into established viral families (n = 3,331). These family-unassigned viruses were grouped at broader taxonomic levels, including phylum-level unclassified groups (111), phylum (1,762), class (300), or order (1,158) level. For example, many WWTP RdRps (n = 1350) and reference RdRps could only be classified into *Kitrinoviricota* phylum without further assigned taxonomic information (Supplementary Fig. 2). These *Kitrinoviricota* RdRps were distributed among three RdRp megataxa (2, 7 and 16, Supplementary Fig. 2 and Table 4). Furthermore, one RdRp megataxon predominantly composed of WWTP-derived RdRps, emerged as a sister lineage to *Ellioviricetes* in the phylogenetic tree, potentially representing a new class within the *Negarnaviricota* phylum (Extended Data Fig. 4). Together, these results depict the most complete landscape of phylogenetic and taxonomical diversity of RNA viruses in the global WWTPs and underscored the necessity for establishing new taxonomic frameworks for RNA viruses.

### Disentangling host associations of RNA prokaryotic viruses in the global WWTPs

Through host inference based on the established Orthornaviran taxonomy, prokaryotic RNA viruses from known phage clades accounted for nearly one-third (3,324, 29.1%) of the total RNA vOTUs richness (*Leviviricetes* n = 3,245 and *Cystoviridae* n = 79) (Fig. 3a). The relative abundance patterns of RNA vOTUs with different host ranges varied across sample types and countries, suggesting the dominance of bacteriophages in nearly all sludge samples. (Extended Data Fig. 5 and Supplementary Text 2).

**Fig. 3.**
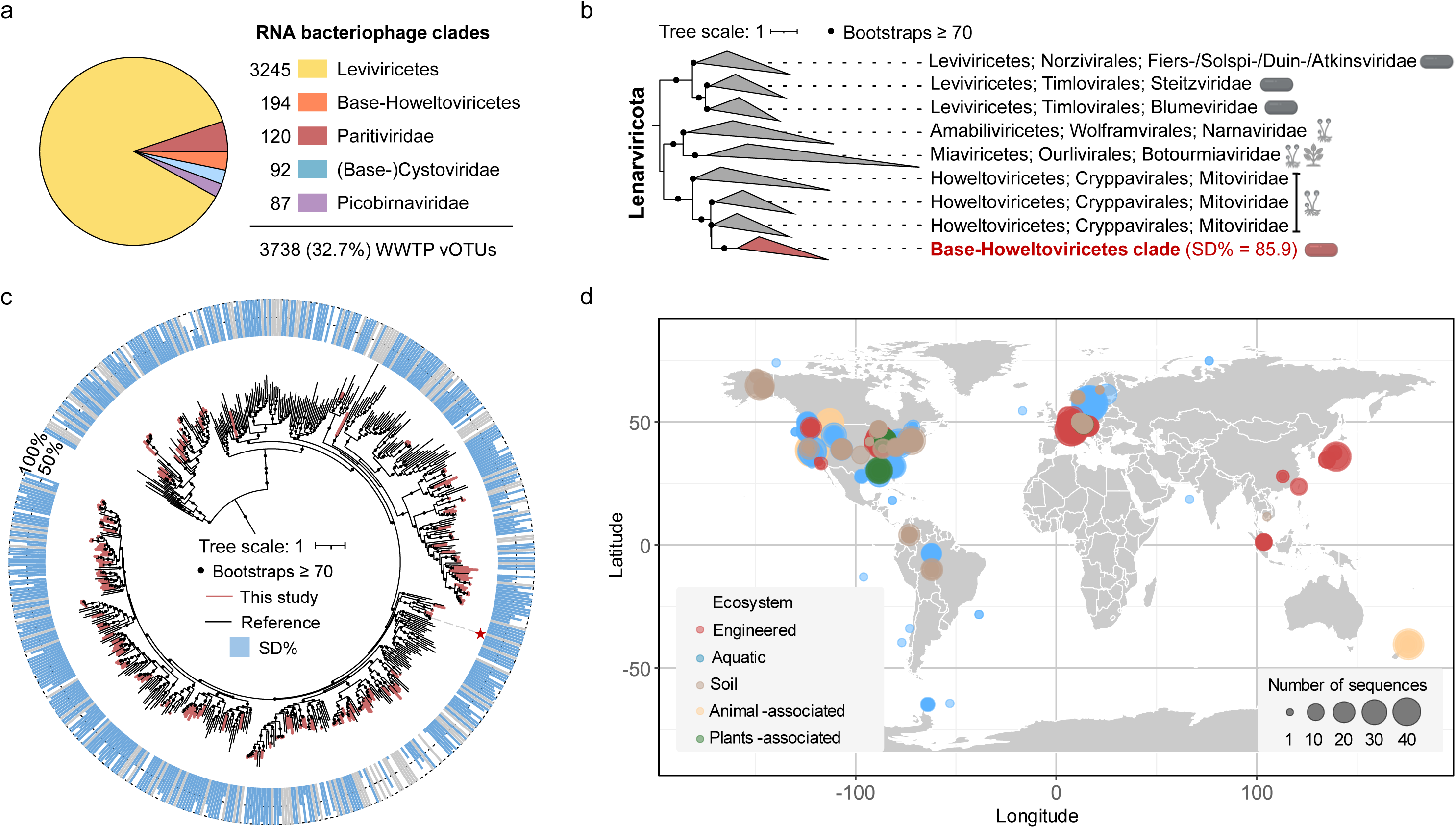
Expanding RNA bacteriophage diversity from global WWTPs. a. The number of WWTP vOTUs from known and novel RNA bacteriophage clades. b. Phylogenetic placement of the base-*Howeltoviricetes* phage clade within the *Lenarviricota* phylum. c. Phylogenetic tree of the base-*Howeltoviricetes* phage clade pruned from the global RdRp tree. Blue outer bars represent the proportion of ORFs associated with the SD motif in the corresponding reference RdRp centroids or vOTUs, while gray bars indicate cases where no ORFs with true start codons were identified, precluding SD motif assessment. The vOTU marked with a star is linked to a probable host based on CRISPR spacer matching. d. Global distribution of the base- *Howeltoviricetes* phage clade. A viral cluster from the GRvOTU set is considered part of the base-*Howeltoviricetes* phage clade if it contains sequence members that belong to the clade’s phylogenetic tree. Sequence members from these clusters are traced back to their corresponding original public samples, and the number of base-*Howeltoviricetes* phage sequences assembled in these samples is counted. The distribution of this phage clade is then visualized according to sample types and geographic locations.

The identification of Shine-Dalgarno (SD) motifs was also used to infer prokaryotic RNA viral clades. Among the RNA vOTUs classified to *Leviviricetes*, 83.3% of the open reading frames (ORFs) were associated with the SD motif. Similarly, 84.9% of ORFs in *Cystoviridae* were linked to the SD motif. Furthermore, several additional clades were discovered to be probable RNA phages due to their enrichment of SD motifs. These included i) *Picobirnaviridae* (n = 87), ii) a subset of *Partitiviridae* (n = 120), iii) a family-unclassified clade basal to *Cystoviridae* (n = 13) (Extended Data Fig. 6), consistent with a recent study^12^, and iv) a clade basal to *Howeltoviricetes* (n = 194, Fig. 3b-c), which was reported here for the first time.

The base-*Howeltoviricetes* phage clade was phylogenetically classified under the family *Mitoviridae* within the *Lenarviricota* phylum (Fig. 3b). Over 85% of the predicted ORFs in this clade were associated with the SD motif (Fig. 3b-c). This generalist phage clade was identified in 390 out of 557 WWTP metatranscriptomes. Additionally, the GRvOTU clustering set facilitated the assessment of the distribution of the base-*Howeltoviricetes* phage clade across public datasets. Based on the phylogenetic tree of this clade (Fig. 3c), all viral sequence members belonging to this clade were extracted from the GRvOTU set. The distribution of this clade was then determined based on the types and geographical locations of the originating samples of the viral sequence members. Base-*Howeltoviricetes* phage clade was found not only present in WWTPs but was also widely distributed across diverse ecosystems globally, including aquatic environments, soil, and samples associated with animals and plants (Fig. 3d).

The CRISPR spacer matching linked one vOTU within the base-*Howeltoviricetes* phage clade to its probable specific host. Several viral contigs from vOTU FD-DS_117671 matched CRISPR spacers identified in bacterial genomes from the NCBI RefSeq database and in contigs from WWTP metagenomic assemblies (Fig. 4a). These matched sequences were taxonomically assigned to the phylum *Campylobacterota*, with the most specific classification identifying *Aliarcobacter cryaerophilus*, a globally emerging foodborne and zoonotic pathogen (Fig. 4a). Virus-host abundance quantification throughout the WWTPs revealed significantly higher relative abundance of both the vOTU representative and its host in wastewater samples compared to sludge samples (Mann-Whitney U test, all *P* < 10^−6^, Fig. 4b). In time-series samples (n = 47, sampled approximately weekly), the abundance of the vOTU representative exhibited a significant positive correlation with the abundance of *Aliarcobacter cryaerophilus* (Spearman r = 0.76, *P* < 10⁻⁹), further supporting the predicted virus-host relationship.

**Fig. 4.**
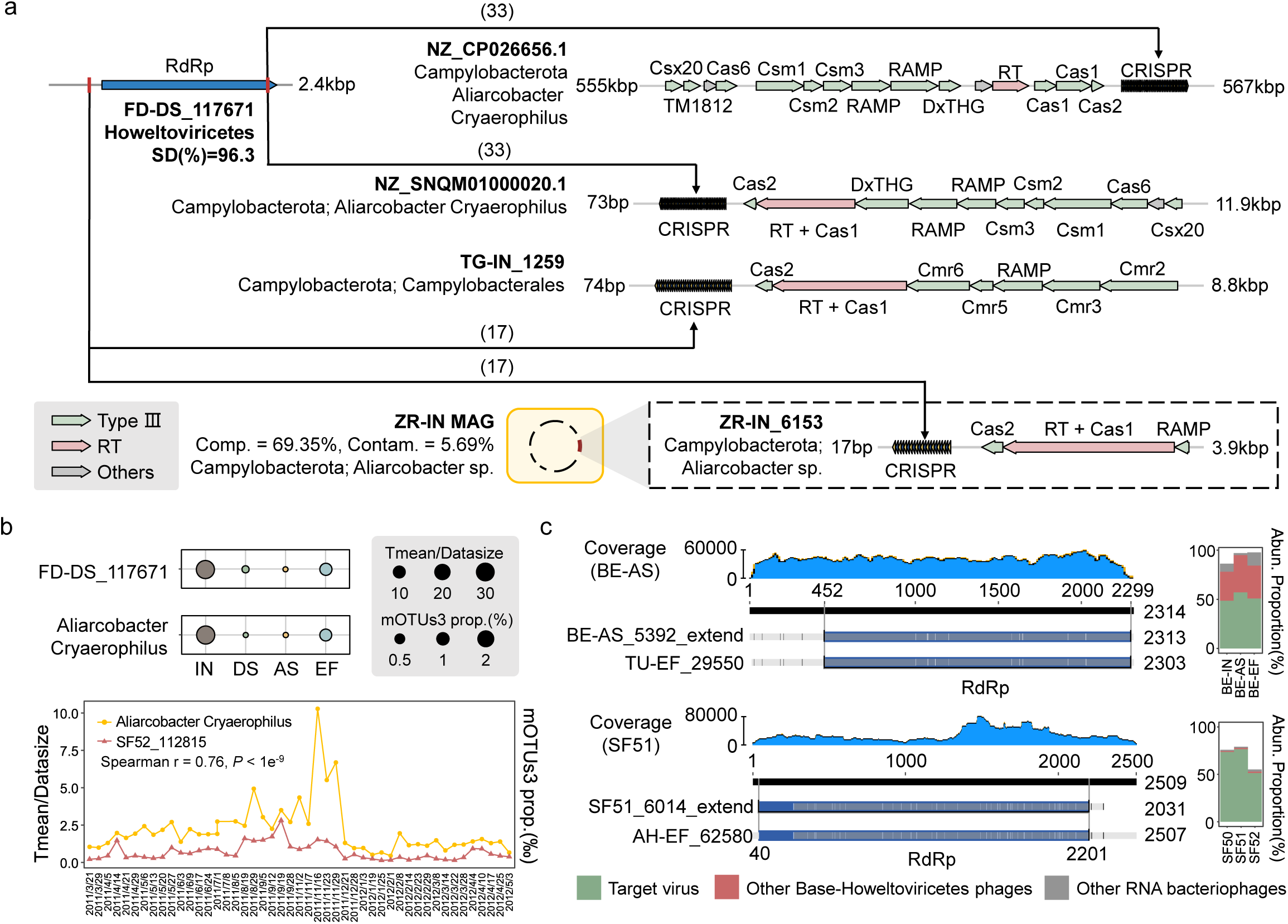
Host association and gene composition of Base-*Howeltoviricetes* phages. a. CRISPR spacer matching links vOTU FD-DS_117671 within the Base-*Howeltoviricetes* phage clade to *Campylobacterota*. Arrows indicate the blastn-short alignments between host spacers and FD-DS_117671, with numbers indicating the count of matching RNA viral contigs within the vOTU. NZ_CP026656.1 and NZ_SNQM01000020.1 are genomes from the NCBI RefSeq database, while TG-IN_1259 and ZR-IN_6153 are WWTP metagenomic contigs. ZR-IN_6153 is binned into a MAG belonging to *Aliarcobacter* sp. Coding genes point right for the (+) sense strand and to the left for the (−) sense strand. b. Relative abundances of FD-DS_117671 and *Aliarcobacter cryaerophilus* during wastewater treatment processes (47 samples from Switzerland, upper panel) and in time-series samples (47 sludge samples from Luxembourg, lower panel). Viral abundance is quantified as the Tmean calculated by CoverM normalized by sample size, and host abundance is estimated using motus3. c. Genome extension for Base-*Howeltoviricetes* phages. Two examples of genome extension are shown. TU-EF_29550 dominates the RNA viral community in BE WWTP samples, with genome extension starting from BE-AS_5392. Similarly, AH-EF_62580 dominates SF50-52 samples, with its genome extension starting from SF51_6014.

In most WWTP samples, *Leviviricetes* was the most abundant group of RNA bacteriophages (Extended Data Fig. 5). However, base-*Howeltoviricetes* phages exhibited substantial abundance in several samples and, in certain cases, became the dominant phage group, contributing to over 50% abundance of the RNA viral community. Examples included TU-EF_29550 in three BE WWTP samples and AH-EF_62580 in SF50-SF52 samples (Fig. 4c). Genome extension of these high-abundance base*-Howeltoviricetes* phages (See Methods) revealed that they likely encode only RdRp (Fig. 4c). Functional annotation did not identify capsid, maturation proteins, or other common RNA viral genes in the genomes of base*-Howeltoviricetes* phages. This observation consisted with the gene composition of their closest phylogenetic relatives, the *Mitoviridae*, a capsid-less viral family that usually encodes only RdRp. Together, these findings provide first insights into the broad distribution, host associations, and genomic features of base*-Howeltoviricetes* phages, a novel RNA bacteriophage clade.

### Prevalence, concentration, and fate of human RNA virus in the global WWTPs

Based on GRvOTU clustering, 21 RNA vOTUs were identified as putative human RNA viruses, including 8 *Mamastrovirus*, 7 *Norovirus* and 3 *Rotavirus* (Fig. 5a). These putative human RNA viruses were all known to potentially cause gastroenteritis in humans. Absolute abundance estimations in the 47 WWTP samples from Switzerland indicated that human RNA vOTUs were significantly reduced during the treatment process, from influent to effluent (Fig. 5b). *Mamastrovirus* and *Rotavirus* vOTUs were consistently detected in nearly all influent and sludge samples in Switzerland, with an absolute abundance peak of 2.01 × 10¹⁰ copies/liter (AD-IN_141867 in TG-AS) for *Mamastrovirus* and 8.42 × 10⁸ copies/liter (FR-INF_306656 in AH-DS) for *Rotavirus*. While the absolute abundance of *Mamastrovirus* vOTUs did not differ significantly between influent and sludge, *Rotavirus* vOTUs exhibited significantly higher concentrations in sludge (Mann-Whitney U test, *P* < 0.05). In contrast, these two viral species exhibited lower prevalence in 277 influent samples from the United States, with *Mamastrovirus* detected in 34 samples and *Rotavirus* in only 5 samples. Norwalk virus was detected in approximately one-third of influent samples in both Switzerland and the United States.

**Fig. 5.**
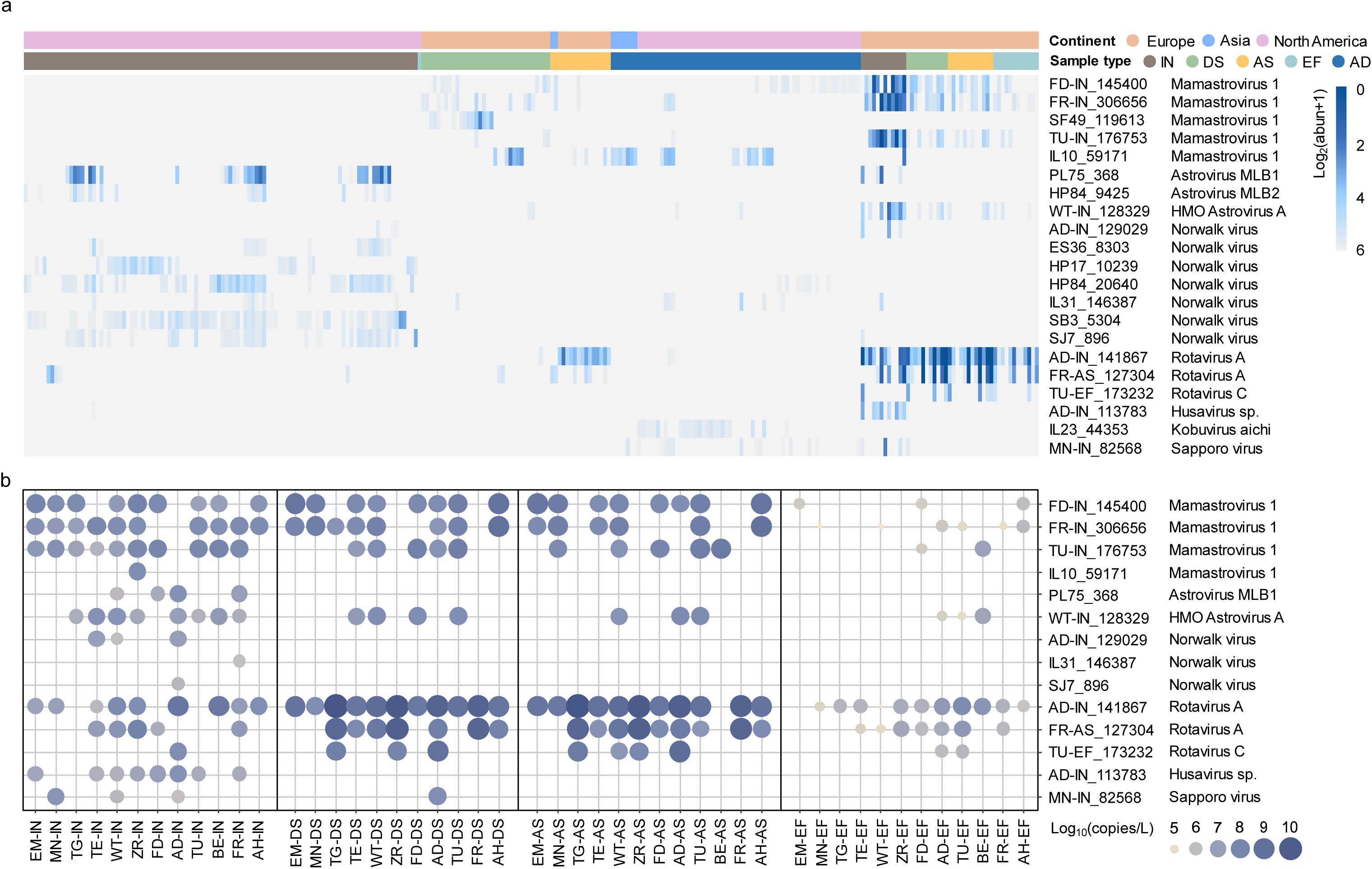
Distribution and absolute concentration of human RNA vOTUs in the WWTPs. a. Heatmap depicting the abundance distribution of 21 human RNA vOTUs identified by species-level clustering. Each row represents a human RNA vOTU and each column represents a sample. Upper annotation columns illustrate the sample location and type. b. Bubble plot showing the estimated absolute concentration of human RNA vOTUs (in copies per liter) in the 47 samples from Switzerland which spiked with mRNA internal standards.

RdRp-based phylogeny was also employed to identify potential human RNA viruses. In addition to the human RNA vOTUs identified based on genome homology match, three novel potential human RNA viruses were discovered through phylogenetic analysis (Extended Data Fig. 7). IL13_9334 and KB7_72351 appeared to be potential novel species within the *Husavirus* genus. Meanwhile, the RdRp of TU-EFF_201332 was positioned between two *Nodaviridae* proteins (Extended Data Fig. 7), which were recovered from human plasma samples and have been identified as potential zoonotic viruses^32^.

To detect human RNA viruses more sensitively, we searched clean RNA reads from metatranscriptomes against the NCBI Virus Database with several evaluation steps (See Methods). A total of 50 human RNA viral species were identified in the global WWTP metatranscriptomes (Extended Data Fig. 8). SARS-COVID-2 was detected in 151 (54.5%) influent samples collected between July 2020 and August 2021 from USA. Five other *Coronaviridae* viral species were detected, including *Alphacoronavirus* 1, *Betacoronavirus*, Human coronavirus HKU15, 229E, and NL63 (Extended Data Fig. 8). Influenza and Hepatitis viruses were also detected in some samples (Extended Data Fig. 8). Reads-based diversity analysis separated the community structure of human RNA viruses in wastewater influent samples from Switzerland and USA (Supplementary Fig. 3), revealing the geographical difference of prevalent viruses and possibly health status between the corresponding populations.

### Ecological roles of RNA viruses in the WWTPs revealed by auxiliary metabolism

To explore the auxiliary metabolism capacities of RNA viruses, a total of 160 RNA viral contig-encoding genes were identified as auxiliary metabolic genes (AMGs) through a series of bioinformatics procedures including gene annotation, authenticity evaluation, and 3D protein structure annotation (Supplementary Table 6). The most prevalent functional types of AMGs were peptidases or proteases (n = 28), followed by ATP-binding proteins (n = 24). Both prokaryotic (n = 9) and eukaryotic (n = 7) ribosomal proteins (RPs) were identified in RNA viral genomes. For instance, AH-IN_3822, which encoded a 60S RP, was classified within *Botourmiaviridae*, a family known to infect fungi or plants (Fig. 6). Phylogenetic analysis of this 60S RP suggested that it was likely acquired from fungal *Mucoromycota* during infection (Supplementary Fig. 4). WT-DS_133589, which encoded a 40S ribosomal protein, was associated with the new megataxon (Fig. 6 and Supplementary Fig. 5). BE-IN_10020 and AD-IN_243634, both from the bacteriophage family *Fiersviridae*, encoded 50S and 30S RPs, respectively (Fig. 6). Phylogenetic analyses indicated that BE-IN_10020 may acquire 50S RP from *Faecalibacterium* within the phylum *Bacillota*, while AD-IN_243634 may acquire 30S RP from *Chitinibacteraceae* within the phylum *Pseudomonadota* (Supplementary Fig. 6-7). Five AMGs related to cellular respiration were identified in the RNA viral genomes, including two glyceraldehyde-3-phosphate dehydrogenase, one citrate synthase, one cytochrome C551, and one ribose-phosphate pyrophosphokinase. One glyceraldehyde-3-phosphate dehydrogenase, involved in glycolysis, was encoded by IL33_2153 from bacteriophage family *Atkinsviridae*, while the citrate synthase gene, involved in the TCA cycle, was encoded by MN-EF_134878 from bacteriophage family *Steitzviridae*, implying a prokaryotic origin for these viral genes (Fig. 6). Additionally, AMGs involved in various other metabolic pathways were also identified, including those related to nucleotide metabolism (n = 9), RNA modification (n = 7), amino acid metabolism (n = 7), and the metabolism of cofactors and vitamins (n = 6).

**Fig. 6.**
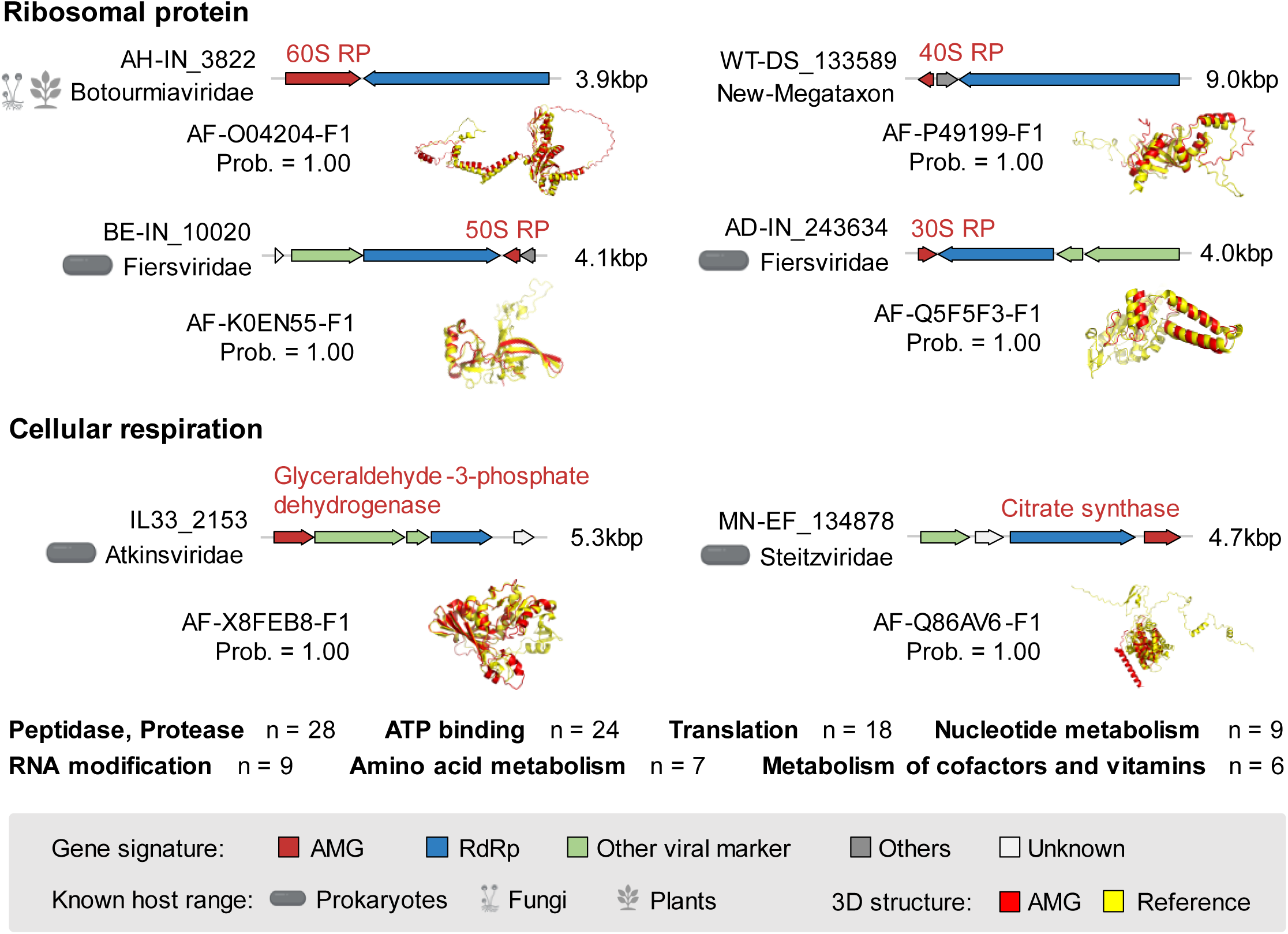
The AMGs encoded by RNA viral genomes revealed the involvement of RNA viruses in various metabolic processes. AMGs are highlighted in red. Coding genes point right for the (+) sense strand and to the left for the (−) sense strand. Below the gene flow, the 3D structures of the AMGs predicted by ESMfold and annotated by Foldseek are displayed. Known hosts are indicated alongside the affiliated taxonomy.

Compared with DNA viruses, interpreting the expression activity of RNA viral genes in metatranscriptomes is inherently challenging because RNA viral genomes and transcripts were interwoven. Here, the abundance comparison between AMG and RdRp within the viral genome in their originating sample was used to indicate the potential expression of AMG. An AMG was considered potentially active if its relative abundance (Tmean) exceeded that of the RdRp gene by more than fivefold (Tmean(AMG) – Tmean(RdRp) > 5×). A total of 19 AMGs were inferred to be potentially expressed in their respective originating samples (Supplementary Table 7). The expressed AMGs included three RPs, two ribonucleases, one elongation factor, and one RNA-binding protein, whose potential expression implied that their hosting RNA viruses may utilize AMGs to enhance transcription, translation, and RNA processing (Supplementary Table 7). Notably, nine unique M15 family peptidases were identified in WWTP RNA viral genomes, seven of which exhibited potential expression activity (Supplementary Table 7). The viruses encoding M15 family peptidases were all derived from the base-*Cystoviridae* clade, a novel bacteriophage group identified through SD motif analysis. In summary, the identified AMGs and their expression patterns point to the dynamic role of RNA viruses in manipulating host cellular machinery and imply multiple virus-host interactions.

## Discussion

This study represents a significant advancement in understanding RNA viruses within wastewater treatment plants (WWTPs), offering the largest and most comprehensive catalog of RNA viruses in these human-essential and environment-friendly engineered microbial ecosystems to date. By creating the first WWTP RNA Viruses Database (WRVirDB), we expand the known diversity of RNA viruses in WWTPs by 67%. This service-oriented resource not only provides a crucial platform for future research into the diversity, ecology, and applications of RNA viruses in WWTPs, but also has profound implications for public health monitoring, particularly in the emerging field of WBE, which promises to improve disease surveillance and public health strategies globally.

### Knowledge Advancements in RNA Virus Diversity and Function

Our metatranscriptomic compendium has refined the global RNA-dependent RNA polymerase (RdRp) phylogenetic tree (Supplementary Text 3), uncovering previously unidentified RNA viral clades, with a particular focus on RNA bacteriophages. These findings demonstrate that WWTPs harbor a remarkably diverse and abundant population of prokaryotic RNA viruses, which are likely integral to shaping microbial communities and biogeochemical processes within these systems (Fig. 2a and Extended Data Fig. 5). Given that that bacteria dominate and drive the microbial ecology of WWTPs^19, 20^, it is not surprising that RNA phages are particularly abundant in these environments. Moreover, the widespread presence of RNA phages in engineered systems and soil environments (Supplementary Fig. 8) is likely influenced by the composition of host organisms, underscoring WWTPs as pivotal ecosystems for uncovering RNA phage and virome diversity.

Our identification of the phage clade base-*Howeltoviricetes*, a group widely distributed across various environments, highlights the underappreciated ecological roles and diversity of RNA phages in WWTPs. Additionally, we reveal that RNA viruses carry a wide array of AMGs of both prokaryotic and eukaryotic origin (Supplementary Table 6). These genes, likely acquired through horizontal gene transfer during viral infection, suggest that RNA viruses are not merely bystanders but participants in influencing host metabolic pathways, energy flow, and element cycling within WWTPs. Such interactions point to the potential for intricate virus-host relationships.

### Implications for Wastewater-Based Surveillance

This study also highlights the potential of metatranscriptomics for wastewater-based surveillance of human viruses, an emerging and promising tool for monitoring infectious diseases ^25, 33^. Metatranscriptomics allows for the simultaneous high-throughput detection of a broad spectrum of RNA viruses, including novel species, thus expanding the database of known human RNA virus (Fig. 5 and Extended Data Fig. 7-8). The detection of SARS-CoV-2 and other human RNA viruses, such as those responsible for gastroenteritis, influenza, and hepatitis, demonstrates the capacity of this broad-spectrum omics approach for virus detection at the read and/or genome levels. Importantly, absolute quantification of human viral loads in WWTPs improves the accuracy of viral load trend assessments (compared with current imperfect relative abundance profiling), enhances the reliability of health assessments (Fig. 5b)^31^. As future quantitative spatial–temporal viral genomic analyses of wastewater samples become more refined, such surveillance could provide both real-time insights into population health status and early warnings for infectious diseases at local, national, and global levels.

### Research Significance and Concluding Remark

This comprehensive study of RNA viruses in WWTPs achieves several major breakthroughs, including the expansion of RNA viral diversity and functionality in engineering systems, and the elucidation of human RNA virus distribution patterns in global wastewater samples. However, the global diversity of RNA viruses in WWTPs remains an open area for further exploration. Some limitations of this study and other RNA virome studies include the reliance on RdRp gene as a universal marker, potentially missing non-RdRp coding fragments of RNA viruses, and the inherent limitations of current sequencing and bioinformatic methodologies. Despite these challenges, the genomic and functional data generated represents a valuable resource for future research and applications.

In conclusion, this study underscores the ecological and public health significance of RNA viruses in WWTPs. By illustrating their roles as key ecological drivers and health indicators, we position RNA viruses as critical components of wastewater ecosystems and other engineered environments. This work contributes to advancing microbial ecology, enhancing public health, and improving disease surveillance strategies, with potential to inform both local and global responses to emerging infectious diseases.

## Methods

### Metatranscriptome collection

To build the first catalogue of RNA viruses of global WWTPs, a total of 557 available metatranscriptomes of WWTP samples were downloaded from the public databases including Integrated Microbial Genomes & Microbiomes (IMG/M), National Center for Biotechnology Information Sequence Read Archive (NCBI SRA), Metagenomic for Rapid Annotations using Subsystems Technology (MG-RAST) and National Genomics Data Center (NGDC) to identify RNA viruses. These metatranscriptomic samples included 289 influent, 80 denitrification sludge, 61 activated sludge, 26 effluent, and 101 anaerobic digester sludge samples from 9 countries (Supplementary Table 1). The 47 samples from Switzerland were sequenced at a similar depth (8.26 ± 1.90 G per sample) and were spiked with mRNA internal standards in the previous study^31^, which enabled the absolute concentration estimation of RNA viruses in these samples. The detailed information for these public metatranscriptomes were listed in Supplementary Table 1.

### Quality control, metatranscriptomic assembly and gene prediction

The raw sequencing reads were trimmed by Trimmomatic (v0.39)^34^ with default parameters. MEGAHIT^35^ has been regarded as a successful assembly tool for metatranscriptomes with efficient memory and computational time^10^. The trimmed reads of each metatranscriptome were assembled into contigs using MEGAHIT (v1.1.3)^35^ with default parameters. Only contigs > l kbp were used for the downstream analysis. ORFs were predicted from assembled contigs using Prodigal (v2.6.3)^36^ with option “-p meta”.

### Identification and evaluation of RNA viruses

The methodology of RNA virus identification referred to a previous study which systemically searched RNA viruses in the global ocean metatranscriptomes^10^. In detail, RNA viruses were identified by iterative HMM searches against virus RdRp HMM profiles (Extended Data Fig. 1). The initial 65 RdRp HMM profiles were downloaded from (doi:10.5281/zenodo.5731488)10. For every iteration, all the protein sequences predicted from the assembled contigs, along with the public virus RdRp protein sequences from NCBI Genbank release 253 (n = 3,694,787), and recent RNA virus studies from ocean (n = 44,828^10^ and n = 4,593^11^) and global metatranscriptomes from diverse ecosystems (n = 329,202)^12^ were searched against the RdRp HMM profiles by hmmsearch (v3.3.2)^37^ with the option “-A”, which supported to trim the portion aligned to the RdRp HMM profile. The hit protein sequences with a bit-score ≥ 30 and with ≥ 70% aligned portions to the best RdRp HMM profiles were retained. The aligned portions of hit protein sequences were clustered into updated RdRp HMM profiles by vFam pipeline (version February 2014)^38^ with default parameters. The RdRp profile HMMs were generated and updated for eight iterations to meet the near saturated number of identified viral RdRps. A total of 56,086 putative virus RdRps and 455 RdRp HMM profiles were identified after eight search-and-updated iterations. The contigs encoded putative RdRps were regarded as putative RNA viral contigs.

To evaluate the authenticity of the putative RNA viruses, false-positive RNA viruses were filtered by 1) a competitive hmmsearch approach^10^ and 2) mapping against corresponding metagenomic datasets. First, 56,086 putative RdRps were searched against the database merged by PfamA (v33.0)^39^ HMM profiles and 455 RdRp HMM profiles constructed in this study. If the putative RdRp showed a best match to a non-RdRp HMM profile, it was considered as a false-positive virus RdRp. All of the 56,086 putative RdRps passed the competitive hmmsearch test, and they all showed a best match to the RdRp HMM profile. Then, based on the hypothesis that RNA viruses (under *Orthornavirae* kingdom) would not be present in the DNA fraction, the occurrence of putative RNA viral contigs in the metagenomes were examined and used to evaluate false-positive RNA viruses. We downloaded additional 134 metagenomes originated from the samples same as part of the WWTP metatranscriptomes. The raw DNA sequences were mapped to the putative RNA viral contigs by Bowtie2 (v2.3.5.1)^40^ in “--very-sensitive” mode. If >20% of the positions in the putative RNA viral contig were mapped by DNA reads in any metagenome, this RNA viral contig was considered as a false-positive RNA virus. In total, 165 putative RNA viral contigs were identified as false-positive RNA viruses and were discarded in this step.

To examine the completeness of virus RdRp domains, RNA viral contigs were translated into protein sequences by transeq (EMBOSS:6.6.0.0)^41^ with all six frames and the standard translational code. The translated protein sequences were searched against the 455 RdRp HMM profiles once more, and the sequences with a bit-score ≥ 30 and with ≥ 90% aligned portions to the best RdRp HMM profiles were considered as RNA viruses containing “complete” (or “near-complete”) RdRp domains. Together, 55,900 RNA viral contigs were identified from 557 WWTP metatranscriptomes, including 44,399 RNA viral contigs contained “complete” RdRp domains.

To check the alternative translation code usage of RNA vOTUs, the ORFs of RNA vOTUs were predicted under all available codes with Prodigal (v2.6.3)^36^. The codon producing the longest average length of ORFs was chosen for the corresponding RNA vOTU.

### RNA virus clustering

#### Clustering of WWTP RNA virus operational taxonomic units

The RNA viral contigs were clustered into RNA vOTUs of 90% ANI across 80% of AF (parameters referred to reference^10^, the scripts used for RNA vOTU clustering were downloaded from https://bitbucket.org/berkeleylab/checkv/src/master/scripts/). The clustered RNA vOTUs were pragmatically at approximately species level^10^. In total, 55,900 RNA viral contigs were clustered into 11,414 RNA vOTUs, including 9,276 RNA vOTU representatives contained “complete” RdRp domains.

#### Clustering of WWTP and publicly known RNA viruses

To evaluate the expansion of RNA viral diversity in this study, RNA viruses identified from the WWTP metatranscriptomes were clustered alongside publicly known RNA viruses at both the genome and RdRp levels. Public RNA viral genome and RdRp sequences were retrieved from NCBI Virus database (accessed on August 13, 2024) and recent studies of RNA viruses in oceanic^10, 11^, soil^13, 18^, and global metatranscriptomes from diverse ecosystems^12^. Based on the global RdRp phylogenetic tree (See section RdRp-based phylogenetic analysis), no RNA vOTUs from WWTPs were assigned to the *Coronaviridae* family. Due to the clustering efficiency considerations, reference sequences of SARS-COVID-2 were excluded from the clustering analysis. Genome-level clustering was performed using the same parameters as for WWTP vOTU clustering (90% ANI and 80% AF). RdRp-level clustering was conducted using MMseqs2^42^ with the parameters ‘--min-seq-id 0.9 -c 0.333 -e 0.1 --cov-mode 1 --cluster-mode 2’^12^. The resulting RNA viral genome clusters and RdRp clusters were termed GRvOTU and RCR90, respectively, and were classified into three categories, ‘novel’, ‘shared’, and ‘reference’, based on the data sources of their member sequences. A viral cluster was considered to be present in WWTP systems if it included members originating from such environments. All public RNA viral genomes were also subjected to genome-level clustering (90% ANI and 80% AF) to establish the background reference for known RNA viral diversity.

#### RdRp-based taxonomy classification

The taxonomic megataxon of RNA vOTUs were determined by a network-based clustering method^10^. The public RdRp reference protein sequences that matched to RdRp HMM profiles after eight hmmsearch iterations were collected and pre-clustered at 50% amino-acid identity by Uclust (v8.0.1623)^43^ with options “usearch – cluster_fast -id 0.5 -sort length”, and resulted in 30,504 public RdRp centroids. Then, 30,504 public RdRp centroids along with 9,276 WWTP RNA vOTUs containing “complete” RdRp domains were merged and used for clustering megataxon of RNA viruses. Pairwise comparisons of these merged RdRp sequences were conducted by BLASTp (v2.11.0) with options “-gapopen 9 -gapextend 1 -word_size 3 -threshold 10”. E-values were extracted and negative-log10-transformed by “mcxload” in MCL (v14-137)^44^ with options “--stream-mirror --stream-neg-log10 -stream-tf ‘ceil(200)’ “. Transformed e-values were used for MCL^44^ network clustering with option “-I 0.9” to establish RdRp megataxon (approximately at class to order level). In total, 22 RdRp megataxon were clustered and labeled by the public reference RdRp centroids with known taxonomic classification within the megataxon. The accuracy of the label for each megataxon was calculated as the number of public RdRp centroids affiliated with the taxonomy divided by the total number of known public centroids (those with labels at phylum level at least) in the megataxon. One megataxon (megataxon 22, n = 68) were mainly consisted of RdRps identified in this study (n = 56) with only two known reference RdRps. This megataxon was regarded as novel megataxon and was not labeled (Supplementary Table 4).

### RdRp-based phylogenetic analysis

#### Establishment of the global viral RdRp tree

The public RdRp centroids, WWTP viral RdRps containing “complete” RdRp domains, along with the group II introns and non-LTR retrotransposons were aligned and used to generate the approximate global tree for viral RdRps. In detail, the protein sequences of group II introns and non-LTR retrotransposons were downloaded from Genbank Release 253. The RdRps within megataxon 18 were affiliated with viral families of *Birnaviridae* and *Permutotetraviridae*, which encoded conserve motifs with the order permutation “C-A-B” instead of the canonical permutation “A-B-C”. Such RdRp permutations can make RdRp sequences unalignable with those of the canonical configuration^11^. Therefore, the RdRp sequences of megataxon 18 (n = 388) were excluded from the analysis of global RdRp tree construction and analyzed individually (Supplementary Fig. 6). In total, public RdRp centroid sequences (n = 30,217), group II introns (n = 950) and non-LTR retrotransposons (n = 83), and WWTP RNA vOTUs containing “complete” RdRp domains (except for RdRps of megataxon 18, n = 9,175), were aligned by MUSCLE (v5.1)^45^ under “-super5” mode. The aligned sequences were trimmed by Trimal (v1.4.rev15)^46^ with sites having more than 20% gaps removed. The alignment of trimmed sequences was used to construct an approximate maximum likelihood tree by the FastTree (v2.1.11)^47^ with options “-wag -gamma”. The constructed tree was illustrated in iTOL (v6)^48^. The tree was rerooted between RdRps and RTs. The family-level taxonomic classification for RdRps was assigned based on the placement of the sequences in the tree, requiring the RdRps fell within a clade to be assigned to a specific family. The family-level taxonomy classification should be consistent in both methods of megataxon clustering and phylogenetic analysis.

#### Phylogenetic analysis for established RNA bacteriophage families

To construct phylogenetic trees for known RNA bacteriophage families, RdRp representatives from RCR90 clusters occurred in the WWTP systems were selected for phylogenetic analysis. Sequence alignment was performing by MUSCLE (v5.1)^45^ and phylogenetic analysis was conducted with IQTREE (v2.1.4-beta)^49^ using parameters ‘-bb 1000’. The resulting trees were illustrated in iTOL (v6)^48^.

### Host prediction of RNA vOTUs

Virus-host relationships were predicted using the following approaches. (i) taxonomy-based prior known host range, (ii) CRISPR spacer match, (iii) SD motif prediction, (iv) species-level clustering, (v) RdRp phylogeny, and (vi) Reads-based alignment. The second and third approaches were designed to identify RNA phages, while the fourth, fifth, and sixth approaches aimed to identify human RNA viruses.

For taxonomy-based host range, the taxonomic classification for RNA vOTUs (See RdRp-based taxonomy classification and RdRp-based phylogenetic analysis) were used to retrieve known host range of each family determined by ICTV proposals.

Type-III and Type-VI CRISPR systems can target RNA and appear possibilities to defend against RNA bacteriophages^50^. To identify potential RNA phages by CRISPR spacer match, all RNA viral contigs were compared to prokaryotic CRISPR spacer sequences to predict virus-prokaryote associations. CRISPR spacers were predicted for all > 10 kb reference bacterial and archaeal genomes downloaded from NCBI Refseq database (https://ftp.ncbi.nlm.nih.gov/genomes/refseq/) with CRT (2.0rev0)^51^. We also predicted CRISPR spacers from metagenomic contigs assembled from 134 corresponding WWTP metagenomes. The DNA contigs were assembled by WWTP metagenomes using metaSPAdes (v3.13.0)^52^ with default parameters. The predicted CRISPR spacers were removed if (i) they included low complexity (consisting of 4– 6bp repeat motifs) or ambiguous bases, (ii) they were ≤ 20bp ^27^. To link RNA viral contigs to predicted CRISPR spacers, only BLASTn-short hits with 0 or 1 mismatch over the whole spacer were considered. The existence, type, and location of CRISPR systems were identified in potential host genomes using GenPept files for reference hosts from the NCBI RefSeq database and gene annotations for WWTP metagenomic assemblies. The ORFs from host DNA contigs were predicted by Prodigal (v2.6.3)^36^ and annotated by HHblits (v3.3.0)^53^ with options “-n 1 -e 0.001” using UniRef30_2023_02 database. Only hits with probability >90% were retained. The predicted virus-host pairs were considered if the matched spacer array was next to an RT-encoding Type III or Type VI CRISPR system in the host genome.

To identify potential RNA phages by SD motif prediction, the information of “rbs_motif” for ORFs of RNA vOTUs with the genetic code identified as above (See Identification and evaluation of RNA viruses) was extracted from Prodigal output files. The canonical SD motifs were identified based on the reference list from the previous study^12^. To identify potential novel prokaryotic RNA viral clades, the viral contigs from two public datasets (the Global Ocean Virome^10^ and The RNA Viruses in Metatranscriptomes Discovery Project^12^) were recruited and also checked for the presence of SD motif as described above. The proportion of SD motif-associated ORFs for each leaf in the global RdRp tree was calculated as the number of SD-carrying ORFs divided by the total number of ORFs with a true start (‘‘start_type’’ different from ‘‘Edge’’ in Prodigal output) within the WWTP RNA vOTU (or the public centroid). The phylogenetic trees of potential prokaryotic RNA viral clades indicated by SD motifs were pruned from the global tree.

To identify potential human RNA viruses by species-level clustering, the GRvOTU clustering results (See RNA virus clustering) was used to identify human RNA viruses. If a viral cluster from GRvOTU dataset included at least one known human RNA virus, the WWTP RNA viral contig within this cluster was assigned to the same species.

To identify potential human RNA viruses by RdRp phylogeny, the public human viral RdRps and WWTP RNA vOTUs containing “complete” RdRp domains were aligned and used to generate a phylogenetic tree. Public human viral RdRp domains retrieved from NCBI genbank release 253 (See “Identification and evaluation of RNA viruses”) were clustered at 95% amino-acid identity to reduce sequence redundancy by Uclust before phylogenetic analysis. Subsequent phylogenetic analysis was performed in the same way as the global RdRp tree construction. The potential human RNA RdRp was identified based on the placement of the sequence in the tree, requiring the RdRp fell within a clade to be assigned to a known clade of human RNA viruses.

To identify potential human RNA viruses by reads-based alignment, all the clean RNA reads from 557 WWTP metatranscriptomes were BLASTn against complete nucleotides of human RNA viruses in NCBI Virus database. The reads matched to reference human RNA viral genomes with ≥ 90% identity along ≥ 80% reads length were considered as potential fragments of the related human RNA viral species. The matched RNA reads were then checked against the NCBI nt database using BLASTn. Reads that matched reference cellular organisms (prokaryotes and eukaryotes) with identity≥ 90% along ≥ 80% read length were discarded. The relative abundance of detected human RNA viral genomes was normalized to RPKM (Reads Per Kilobase of transcript per Million mapped reads) based on reference genome length and the size of metatranscriptome. The relative abundance of human RNA viral species was calculated by summing the RPKM value of the viral organisms belonging to the viral species. Only human RNA viral species with a total read count > 10 in the samples were retained for analyses.

A mock metatranscriptomic dataset, composed of Illumina sequencing data from bacterial transcriptomes and human RNA viral genomes, was generated to further assess the feasibility of reads-based detection of human RNA viruses^54^. Eight Illumina transcriptomic datasets from pure-culture bacteria isolated from WWTP samples were downloaded from NCBI SRA Database (Supplementary Table 9). The mock Illumina sequencing data for eight human RNA viruses were generated by Mason (v2.4.0)^55^ with different sequencing depth (from 20 to 40,000 reads). The merged mock metatranscriptome has 172.5 million reads in total with 72.3k simulated human RNA viral reads (Supplementary Table 9). The mock metatranscriptome was also subjected to the same BLASTn alignment steps against the NCBI Virus and NCBI nt databases. The alignment results were used to evaluate the accuracy of reads-based detection of human RNA viruses. In the mock community, 99.99% of reads belonging to the human RNA viral genome were successfully recovered, and only 28 sequences from bacterial transcriptomes were incorrectly recruited (Supplementary Table 9), suggesting the reads-based examination for human RNA viruses from global WWTP metatranscriptomes in this study is confident and accurate.

### Functional gene annotation of RNA viral contigs

The functional annotation of predicted ORFs with the genetic code identified as above (See Identification and evaluation of RNA viruses) was determined by one iteration of HHblits (v3.3.0)^53^ with options “-n 1 -e 0.001” using UniRef30_2023_02 database (https://www.user.gwdg.de/~compbiol/uniclust/2023_02/UniRef30_2023_02_hhsuite.t ar.gz). Only hits with >90% probability score were used for functional annotation. The predicted ORFs were also searched against the NCBI nr database by BLASTp with a bitscore > 50 were retained.

### Identification and phylogenetic analysis of RNA virus-encoded AMGs

The RNA virus-encoded genes were regarded as AMGs if they met the following requirements: (i) both HHblits best hit in the UniRef database and BLASTp best hit in the NCBI nr database were originated from cellular organism. (ii) No DNA reads from the 134 corresponding WWTP metagenomes could be perfectly mapped to the AMG regions of the RNA viral contigs. (iii) The 3D protein structure annotation was consistent with the homology-based annotation. The 3D protein structures of probable AMG sequences were predicted with ESMFold^56^, annotated with Foldseek (9.427df8a)^57^ (probability threshold set to 0.9), and visualized in PyMOL (v3.0.5)^58^.

To investigate the probable evolutionary origin of RNA virus-encoded RPs, sequences similar to four types of RPs were recruited from the NCBI nr database. The viral and reference RP sequences were aligned by MUSCLE (v5.1)^45^ and analyzed by IQTREE (v2.1.4-beta)^49^. Phylogenetic trees were visualized in iTOL (v6)^48^.

### Quantitative profiling and diversity analysis of RNA vOTUs

#### Quantification of RNA vOTUs

To calculate relative abundance of RNA vOTUs in each sample, clean reads from each metatranscriptome were first trimmed off polyA and polyp stretches using bbduk (v37.62) (https://jgi.doe.gov/data-andtools/bbtools/) with options “literal=AAAAAAAAAA,TTTTTTTTTT, hdist=2 k=10 minlength=30, ktrim=r restrictright=20; ktrim=l restrictleft=20” for three times. WWTP RNA viral contigs were also trimmed off polyA and polyT stretches. PolyA/T-trimmed clean reads in each metatranscriptome were individually mapped against polyA/T-trimmed RNA viral contigs using Bowtie2 (v2.3.5.1)^40^ with options “--local -D 20 -R 3 -N 0 -L 16 -i S,1,0.50 --very-sensitive --non-deterministic -I 0 -X 2000”^10^. The relative abundance of RNA vOTUs in each sample was calculated by CoverM (v0.6.1, https://github.com/wwood/CoverM) with options “--min-read-percent-identity 90 -- min-read-aligned-percent 75 -m trimmed_mean --min-covered-fraction 30”, which recruited reads that mapped with ≥ 90% identity ≥ 75% of the read length and restricted RNA viral contigs should be covered ≥ 30% length horizontally by mapped reads. The calculated Tmean abundances for RNA viral contigs were normalized by the number of mapped reads as filtered by CoverM. The final relative abundance of RNA vOTUs were calculated by summing the normalized abundances of contigs belonging to the RNA vOTUs.

#### Quantification of AMGs

To determine the relative abundance of RNA viral ORFs in its originated sample, the polyA/T-trimmed RNA clean reads from each metatranscriptome were mapped to RNA viral ORFs individually using the ‘--very-sensitive’ mode with Bowtie2 (v2.3.5.1)^40^. The relative abundance of each ORF was then calculated as the trimmed mean abundance, determined using CoverM (v0.6.1) with the options “--min-read-percent-identity 100 --min-read-aligned-percent 70 -m trimmed_mean --min-covered-fraction 70”.

Due to the intermingling of RNA viral genomes and transcriptional products in metatranscriptomes, determining whether RNA viral genes are actively expressed is challenging. However, the potential expression activity of AMG can be inferred by comparing the relative abundance with RdRp within its respective viral genome. An AMG was considered potentially active in its originated sample if: Tmean (AMG) – Tmean (RdRp) > 5×. It was important to note that these bioinformatics steps identify AMGs more likely to be actively expressed, though sequencing inaccuracies, sampling size fractions and gene expression patterns (e.g., actively expressed AMGs with lower expression levels than RdRp cannot be identified) might affect the prediction.

### Absolute quantification of RNA vOTUs in mRNA internal standards spiked samples

To estimate absolute abundance of RNA vOTUs in 47 metatranscriptomes from Switzerland, the statistical information of spiked mRNA internal standards as referred to reference^31^ was used in this study. The absolute abundance of RNA vOTU in each sample was measured in both “copies/g-VSS” and “copies/L” rank using the equation 1 and 2, respectively.

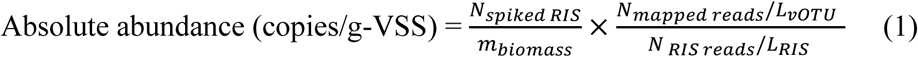

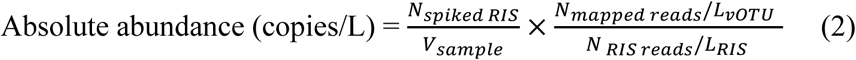

N_spiked RIS_ was the copy number of spiked mRNA internal standards in the sample, m_biomass_ was the mass of collected volatile suspended solids of the sample, N_mapped reads_ was the number of reads mapped to the RNA vOTU in the metatranscriptome, L_vOTU_ was the length of the RNA vOTU, N_RIS reads_ was the number of reads mapped to the mRNA internal standards in the metatranscriptome, L_RIS_ was the length of the mRNA internal standards.

### Diversity analysis of RNA vOTUs

The α-diversity of RNA vOTUs in the samples was measured with the Shannon’s index calculated by function “diversity” in “vegan” package^59^ in R. The β-diversity of RNA vOTUs in the samples was measured using the Bray-Curtis dissimilarity method calculated by function “vegdist” in “vegan” package^59^ in R, using the log-transformed relative abundance table of RNA vOTUs in the samples as input. Then, the Non-Metric Multidimensional Scaling (NMDS) analysis was performed using function “metaMDS” in R package “vegan”^59^. The analysis of Similarity (ANOSIM) was performed using function “anosim” in R package “vegan”. The difference of Shannon index H and absolute concentration of RNA vOTUs between pairwise compartments from Switzerland was determined by the Mann−Whitney U test using the function “wilcox.test” with option “paired = FALSE” in R. Host taxonomy, host-virus abundance correlation and genome contents of base-*Howeltoviricetes* bacteriophage clade

Several viral contigs from vOTU FD-DS_117671, belonging to the base-*Howeltoviricetes* phage clade, matched CRISPR spacer identified in both bacterial genomes from the NCBI RefSeq database and WWTP metagenomic contigs. Two potential host genomes from the NCBI RefSeq database were both classified as *Aliarcobacter Cryaerophilus*. To determine the taxonomic classification of two potential host contigs (TG-IN_1259 and ZR-IN_6153) from WWTP metagenomic assemblies, metagenome-assembled genomes (MAGs) were reconstructed from the respective samples (TG-IN and ZR-IN) using metaBAT2 (v2.12.1)^60^. MAG completeness and contamination were evaluated with CheckM (v1.0.12)^61^, and taxonomy was assigned using GTDB-tk (v2.1.1)^62^. ZR-IN_6153 was successfully binned into a MAG with 69.35% completeness and 5.69% contamination, and classified as *Aliarcobacter sp.*. Since TG-IN_1259 could not be binned into any MAG, its taxonomy was determined using CAT (v5.2.3)^63^.

The viral-host abundance of vOTU FD-DS_117671 was examined across two sample series: i) 47 samples from Switzerland, representing virus-host abundance changes throughout the wastewater treatment process. ii) 47 time-series samples from Luxembourg, capturing the temporal dynamics of virus-host abundance. From the Luxembourg dataset, only the first 47 of 68 samples were included in the correlation analysis because, according to the original metadata, these samples were collected at weekly intervals, ensuring consistent temporal resolution, and sequenced in the same batch, reducing technical variability. The FD-DS_117671 vOTU comprised 53 contigs. As FD-DS_117671 originated from the Swiss sample series, it was designated as the representative sequence for this vOTU in the Swiss dataset. In the Luxembourg sample series, SF52_112815 was selected as the representative sequence for this vOTU, as it was the longest contig recovered from the Luxembourg dataset. To calculate the relative abundance of representative sequence in the related dataset, polyA/T-trimmed clean reads in the corresponding metatranscriptomes were mapped against representative sequence using Bowtie2 (v2.3.5.1)^40^ under ‘--very-sensitive’ mode. The relative abundance of the vOTU representative sequence in each sample was calculated as the trimmed mean abundance, determined using CoverM (v0.6.1, https://github.com/wwood/CoverM) with the options “--min-read-percent-identity 90 --min-read-aligned-percent 75 -m trimmed_mean --min-covered-fraction 30” divided by the size of the metatranscriptome (in Gb). The relative abundance of potential bacterial host (*Aliarcobacter Cryaerophilus*) in the two datasets were determined by mOTUs3^64^ (v3.0.3). Then, the correlation between host and viral abundance was calculated by the Spearman correlation coefficient.

In the BE samples from Bern in Switzerland and the SF50-SF52 samples from Luxembourg, base-*Howeltoviricetes* phage clade showed remarkably high abundance (abundance proportion > 50%). Therefore, near-complete genomes of the highly abundant RNA phages may be reconstructed from these samples. For example, the vOTU TU-EF_29550 from base-*Howeltoviricetes* phage clade accounted for 57.1% of the total RNA vOTU abundance in the BE-AS sample. However, no RNA viral contig members from vOTU TU-EF_29550 were assembled in BE-AS sample. To check this discrepancy, all contigs from the BE-AS sample were aligned to TU-EF_29550 using BLASTn. A total of 83 contigs were identified >95% identity over >95% coverage with TU-EF_29550. These contigs were all less than 1 kb so they were not included in the RNA virus identification step. The fragmented assembly was likely caused by the microdiversity of the RNA virus. Among these 83 contigs, the longest contig (BE-AS_5392, 863bp) was selected as the starting sequence for genome extension. BE-AS_5392 shared 97.7% sequence identity with TU-EF_29550 over the full length. Based on genome extension method described in the reference^65^, BE-AS_5392 was extended to a final length of 2,313 bp. Similarly, in the sample SF51 from Luxembourg, the vOTU AH-EF_62580 from base-*Howeltoviricetes* phage clade accounted for 76.6% of the total RNA vOTU abundance, and the related contig SF51_6014 was extended to a final length of 2,031bp in the SF51 sample.

## Data availability

The sequences of all RNA vOTUs and RdRps identified in this study and the detail information of RNA vOTUs were available at https://github.com/emblab-westlake/WRVirDB. The accession number in a publicly available database will be provided during manuscript submission and upon request.

## Acknowledgements

This work was supported by the Zhejiang Provincial Natural Science Foundation of China under (Grant No. LR22D010001), the “Pioneer” and “Leading Goose” R&D Program of Zhejiang (2024SSYS0032), and the Westlake University-Muyuan Joint Research Institute (WU2024MY004) and the HRHI program 202309010 of Westlake Laboratory of Life Sciences and Biomedicine at Westlake University. The authors thank Ms. Yisong Xu, Ms. Jiajing Guo, Mr. Guoqing Zhang and Ms. Xinyu Huang at Westlake University for their helpful suggestions and technical support. We thank the Westlake University HPC Center for computation support.

## Author contributions

F.J. conceived the project, obtained funding, and supervised the study. F.J. and L.Y. wrote the manuscript. L.Y. performed bioinformatics analysis, statistical analysis, and data visualization. L.C refined viral genomics analysis and data visualization. H.Y., J.Z. and R.S revised the manuscript and provided valuable suggestions for its improvement. F.J. finalized the manuscript and all authors approved the final version.

---

**Extended Data Fig. 1.**
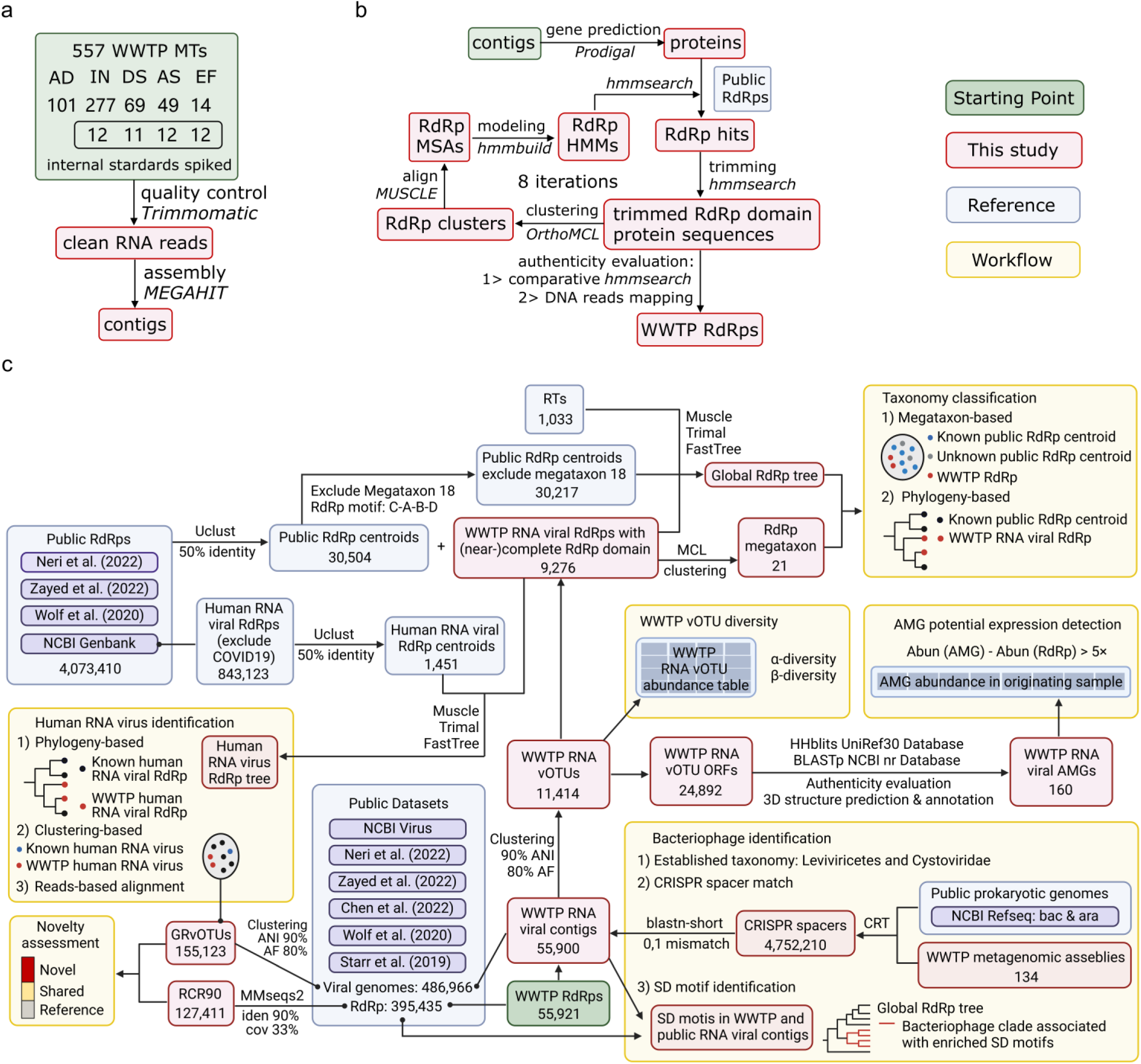
The bioinformatic pipeline for studying RNA viruses in the WWTPs in this study. a. A total of 557 metatranscriptomes were downloaded from public databases and used for RNA virus exploration. Among the collected metatranscriptomes, 47 samples from Switzerland were spiked with mRNA internal standards enabling the estimation of the absolute concentration of RNA viruses in these samples. The metatranscriptomes were first quality controlled and then assembled into contigs. b. After gene prediction, putative viral RdRps were identified by an iterative hmmsearch approach. The authenticity of identified RdRps was assessed by a comparative hmmsearch of putative viral RdRps against RdRp and non-RdRp profiles as well as checking the occurrence of putative viral RdRp-encoding sequences in the metagenomes. c. The RNA vOTUs were clustered at 90% average nucleotide identity (ANI) over 80% alignment fraction of the shorter sequence (AF). Then, to systemically investigate the diversity and ecological roles of RNA viruses in the WWTPs, bioinformatic analyses including taxonomic and phylogenetic classification, RNA bacteriophage identification, human RNA virus identification, and AMG annotation and expression detection were conducted for WWTP RNA viruses.

**Extended Data Fig. 2.**
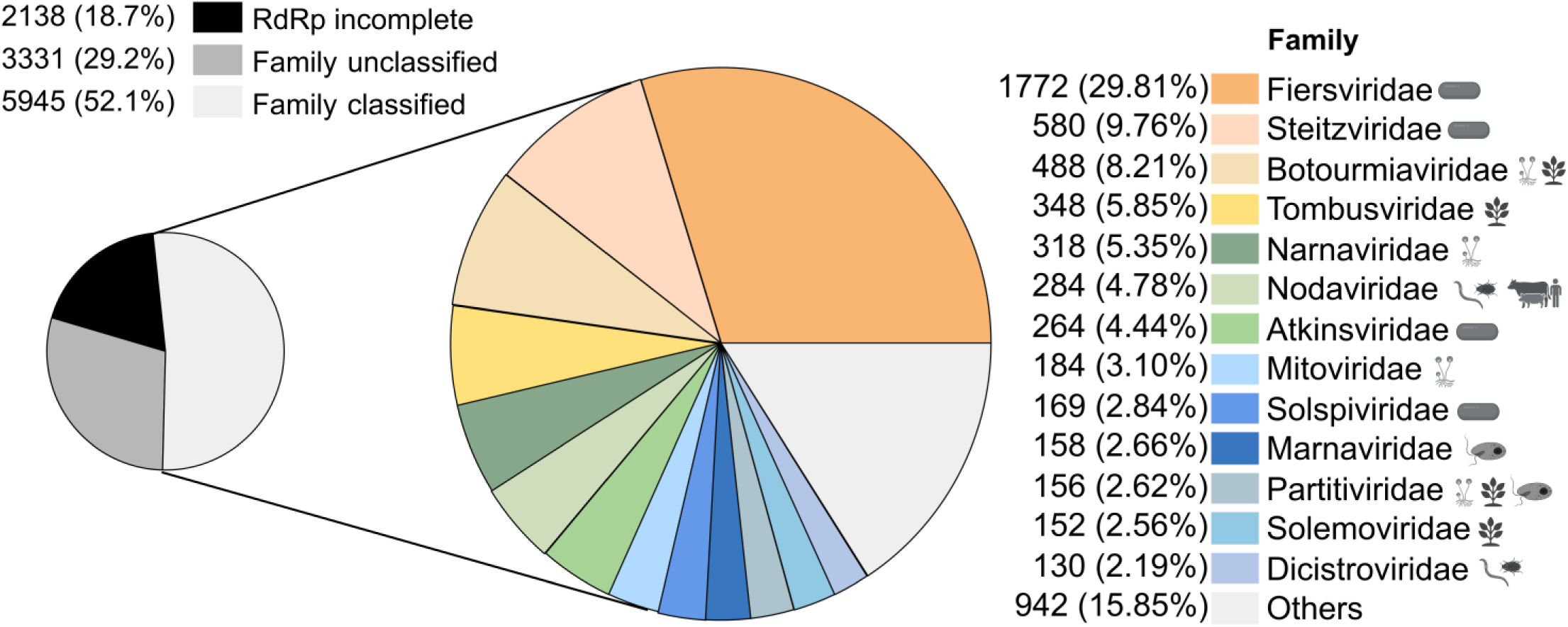
Taxonomy-based richness of RNA viruses in the WWTPs. The richness percentage of RNA vOTUs at the family level. Among 9,276 RNA vOTUs encoding “complete” RdRp domains, 5,945 RNA vOTUs were classified into established viral families. Host range icons next to family names represent their known host ranges.

**Extended Data Fig. 3.**
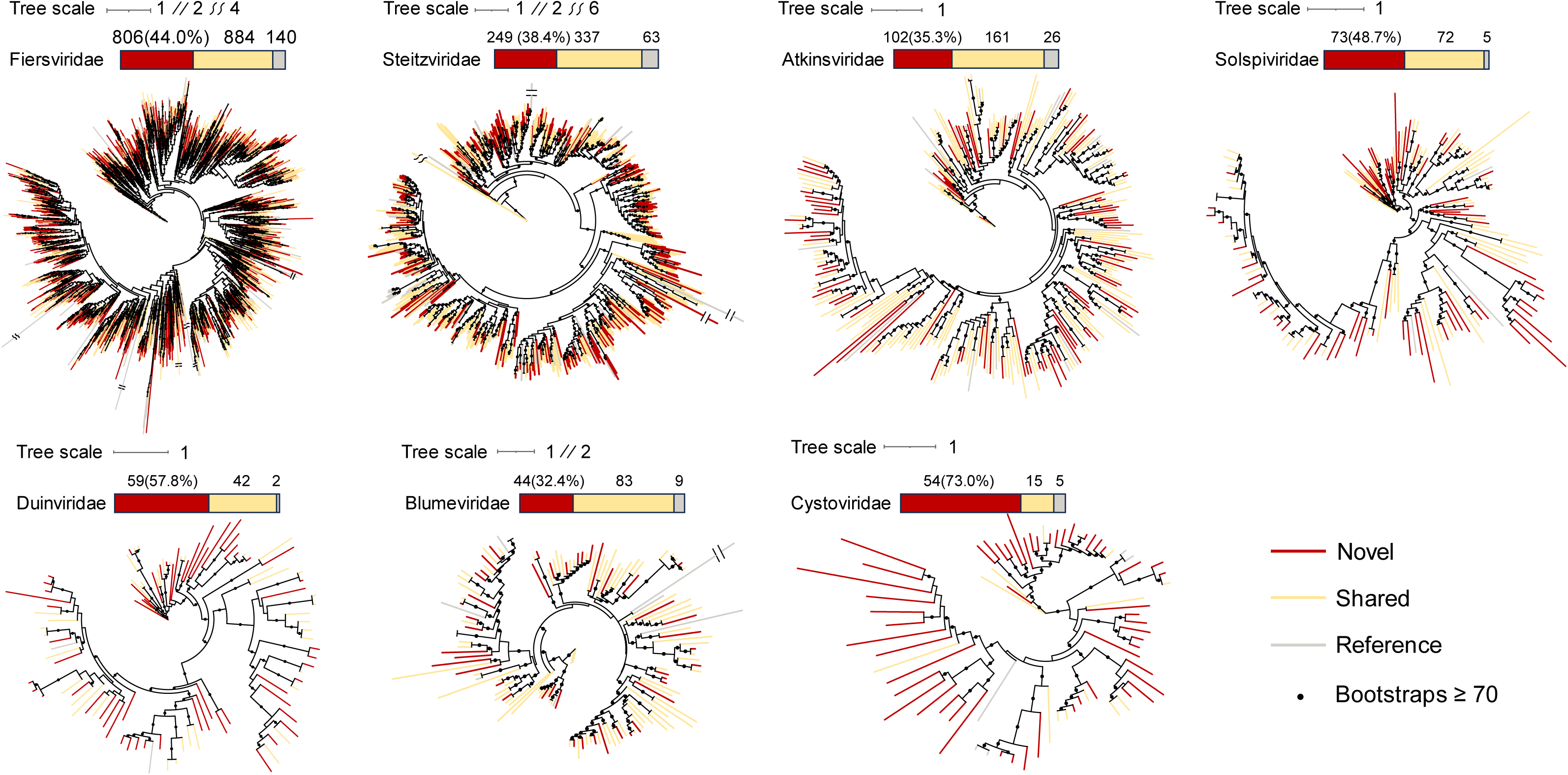
Phylogenetic trees of established RNA bacteriophage families. Each tree includes RdRps from RCR90 set occurred in at least one engineering sample. Leaf colors indicate the type of the corresponding RCR90 RdRp cluster as either “novel,” “shared,” or “reference.“

**Extended Data Fig. 4.**
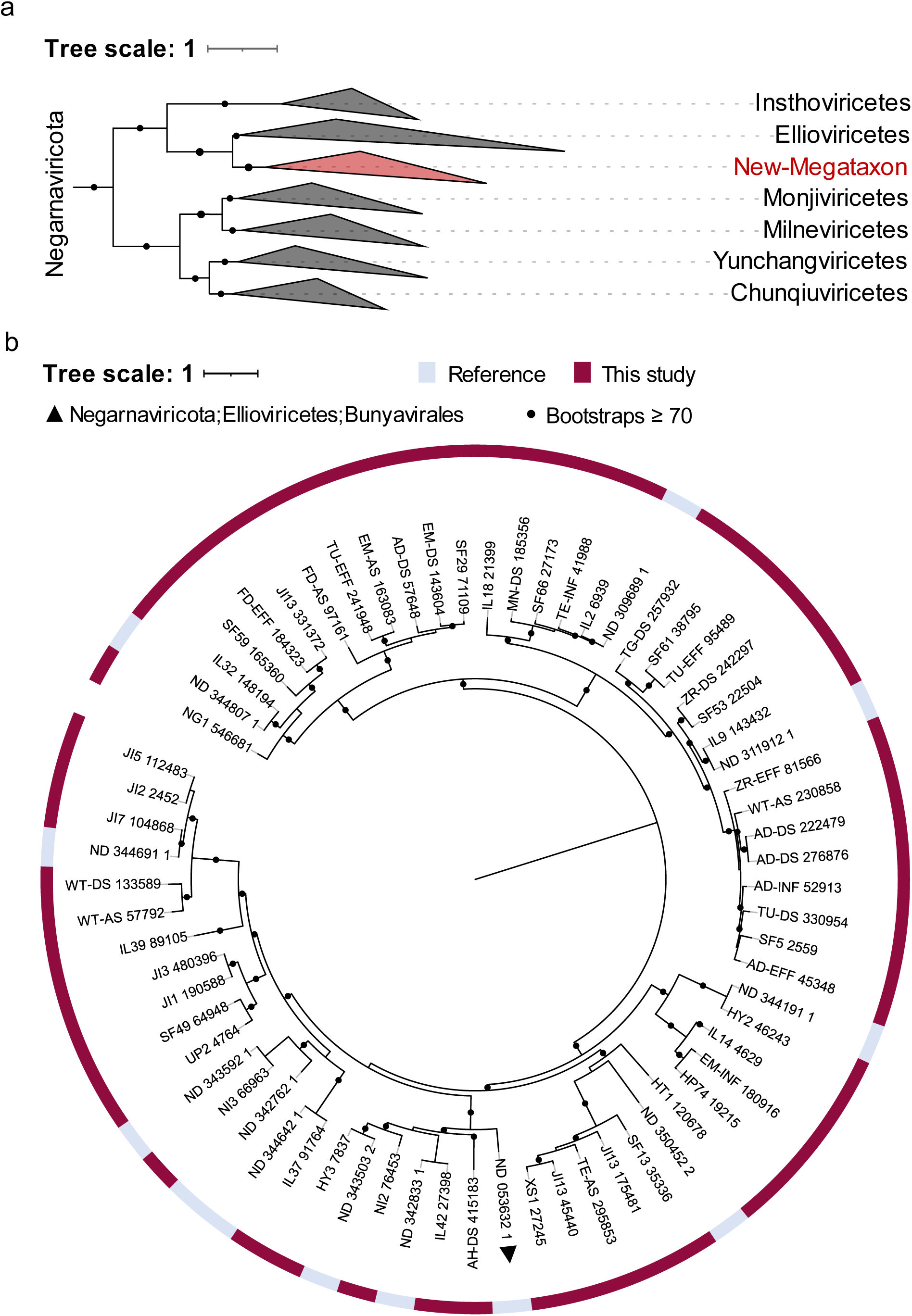
RdRp phylogenetic tree of the newly identified megataxon. This megataxon primarily comprises RdRps from the WWTPs and unclassified reference RdRp centroids. The outer color strip indicates the origin of the RdRp sequences (either from this study or from reference datasets). Among the reference sequences, only ND_053632_1 is annotated with a taxonomic label (*Bunyavirales*), whereas the remaining reference sequences lack taxonomic classification.

**Extended Data Fig. 5.**
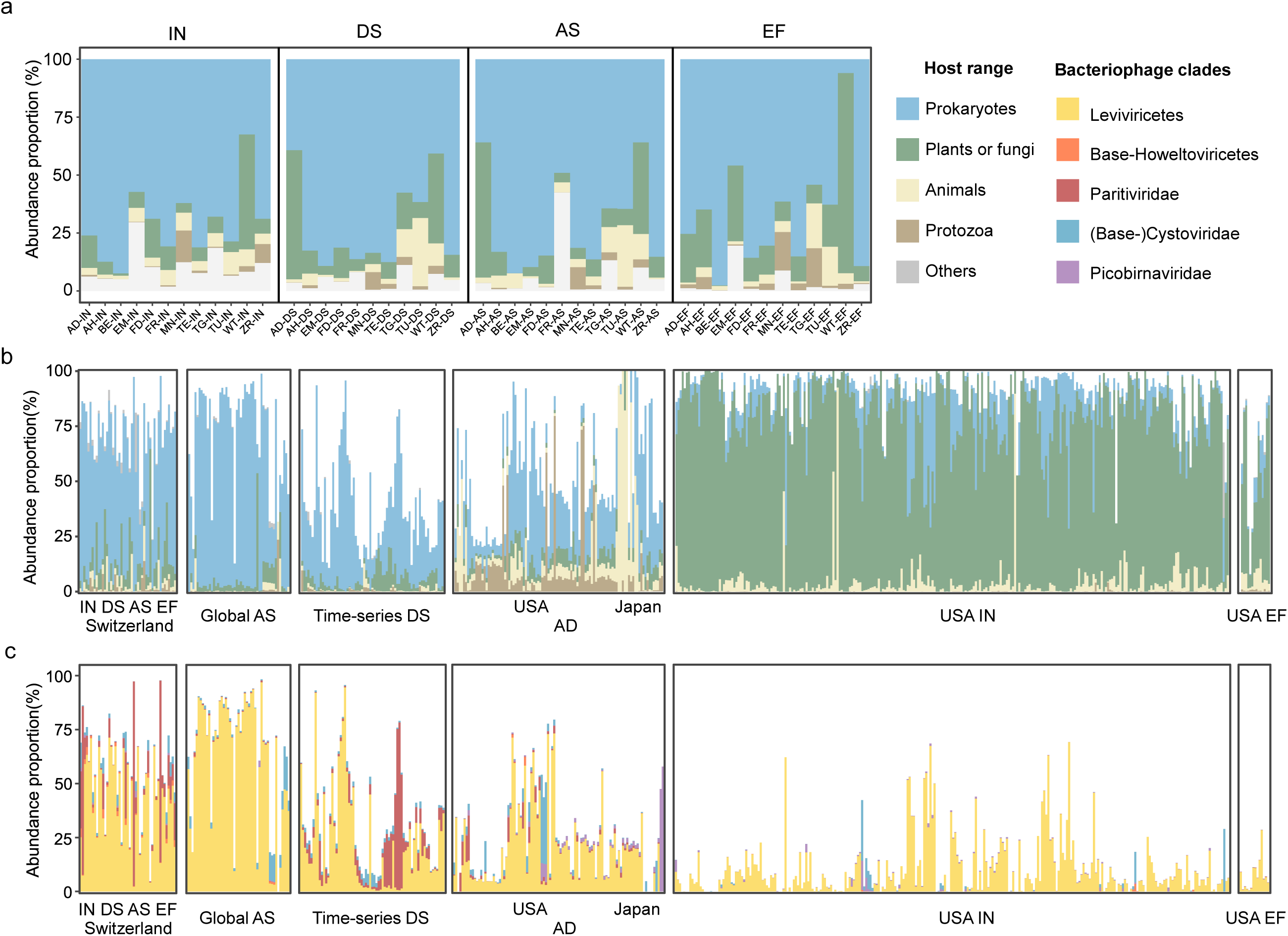
The abundance proportion of RNA viruses in the WWTPs. The abundance proportion of RNA vOTUs in the samples from Switzerland (a) and all samples (b) grouped by their host ranges suggested by affiliated taxonomy. The abundance proportion of RNA bacteriophage clades in all samples (c). The host range of each RNA viral lineage was determined based on: 1) ICTV proposals and 2) previously unknown bacteriophage clades identified in this study (*Picobirnaviridae*, part of *Partitiviridae*, Base-*Cystoviridae*, and part of Base-*Howeltoviricetes*).

**Extended Data Fig. 6.**
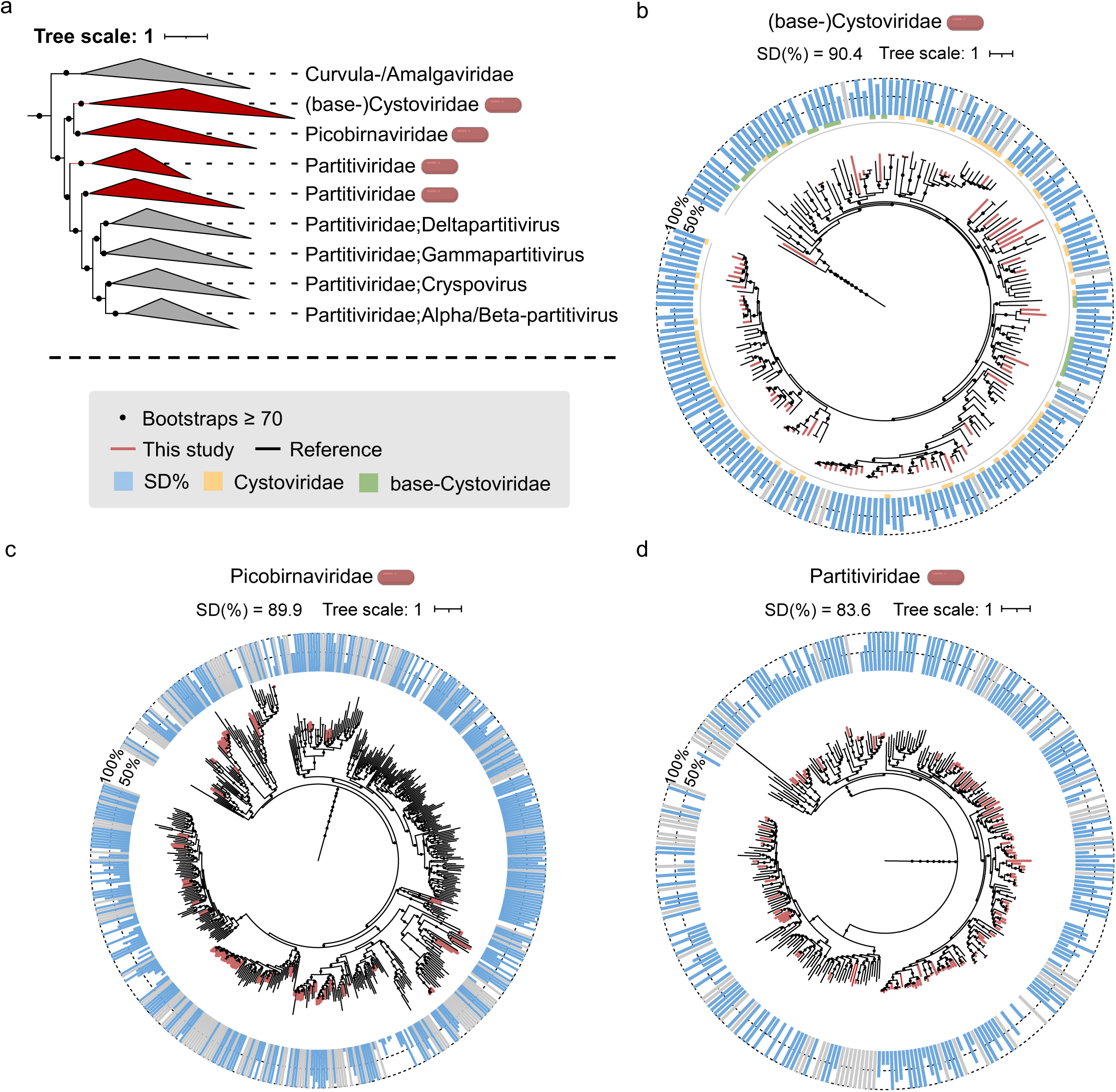
RNA bacteriophage clades in phylum *Pisuviricota* identified through Shine-Dalgarno motifs. a. Phylogenetic placement of RNA bacteriophage clades within the *Pisuviricota* phylum. All RNA viral families in the tree are assigned to the *Durnavirales* order, except for (base-)*Cystoviridae*, which is currently assigned to *Duplornaviricota* by ICTV but was placed under *Pisuviricota* in both this study and a recent study (Neri et al.). b–d. Phylogenetic trees of (base-)*Cystoviridae* (b), *Picobirnaviridae*, and a subset of *Partitiviridae* (d) were pruned from the global RdRp tree. Blue outer bars represent the proportion of ORFs associated with the SD motif in the corresponding reference RdRp centroids or vOTUs, while gray bars indicate cases where no ORFs with true start codons were identified, precluding SD motif assessment. The RdRps of *Cystoviridae* and base-*Cystoviridae* are phylogenetically intermingled. Thus, their phylogeny is displayed together in Figure b, with an additional outer color strip indicating their corresponding taxonomy.

**Extended Data Fig. 7.**
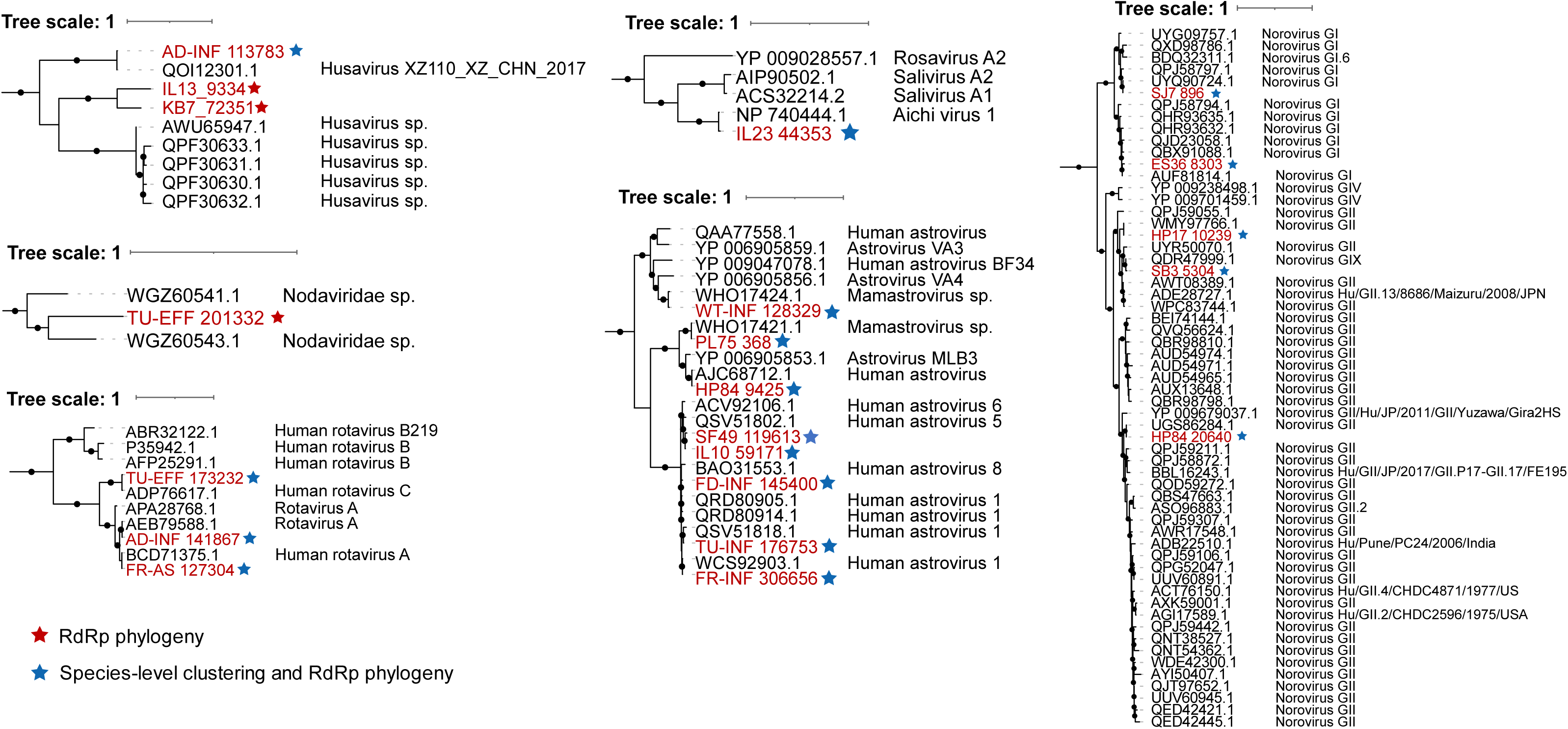
Potential human RNA viruses identified through RdRp phylogeny. RdRps labeled in red represent sequences from WWTP RNA vOTUs, while the remaining sequences are reference RdRp domain sequences obtained from the NCBI Genbank Database. Human RNA viruses identified through both species-level clustering (See Methods) and RdRp phylogeny are marked with a blue star. Human RNA viruses identified exclusively through RdRp phylogeny are marked with a red star and are considered novel human RNA viruses, at least at the species level.

**Extended Data Fig. 8.**
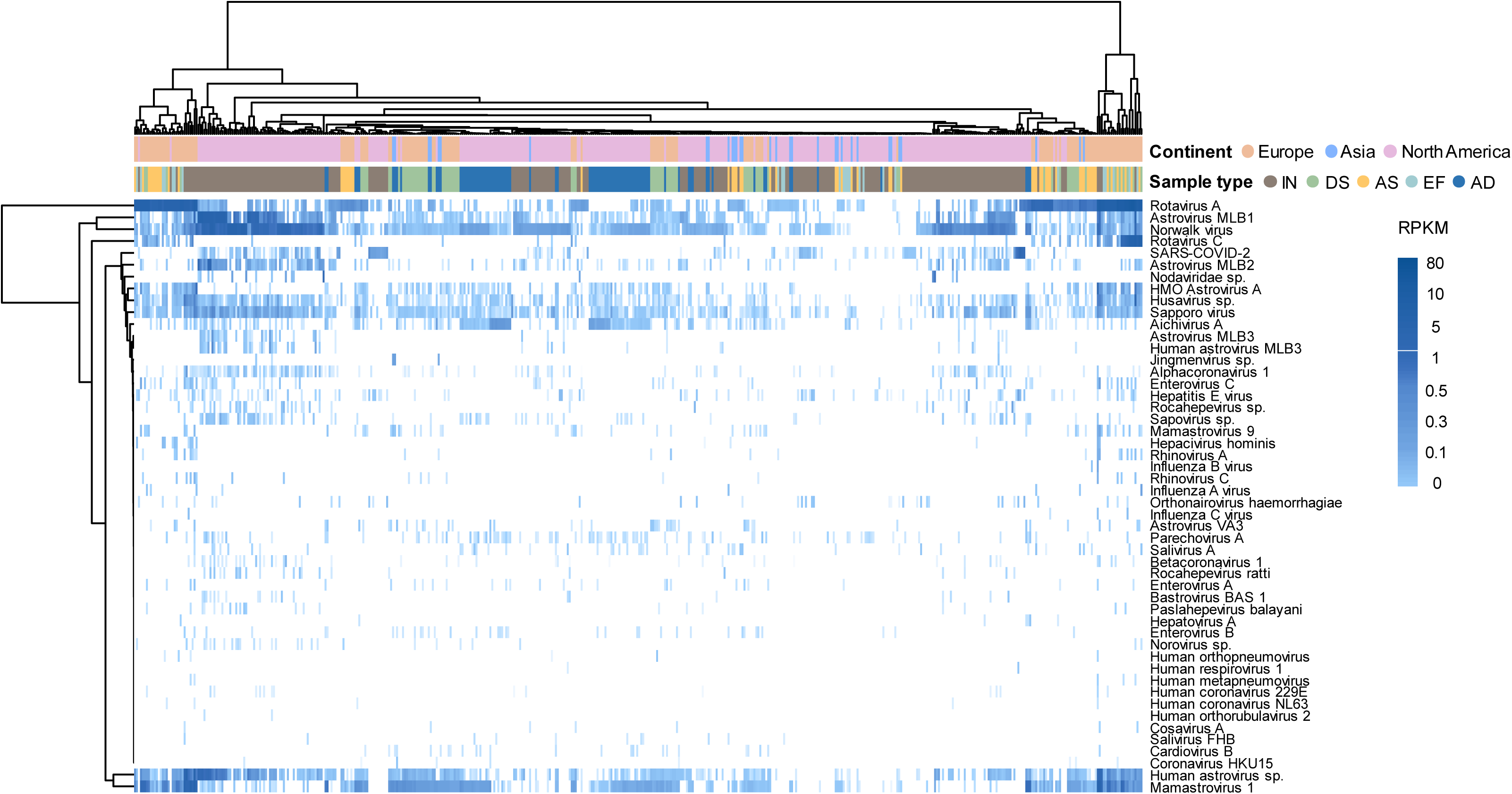
Detection of potential human RNA viruses through short-read alignments. The heatmap displays the RPKM values of human RNA viruses identified through read-based alignment. The alignment was performed using blastn to align clean RNA reads from the metatranscriptomes against the NCBI Virus database, with a 90% identity threshold and 80% read length coverage as cutoffs. A constructed mock community validated that the read-based approach for detecting human RNA viruses in global WWTP metatranscriptomes is reliable (See Methods).

**Supplementary Fig. 1.**
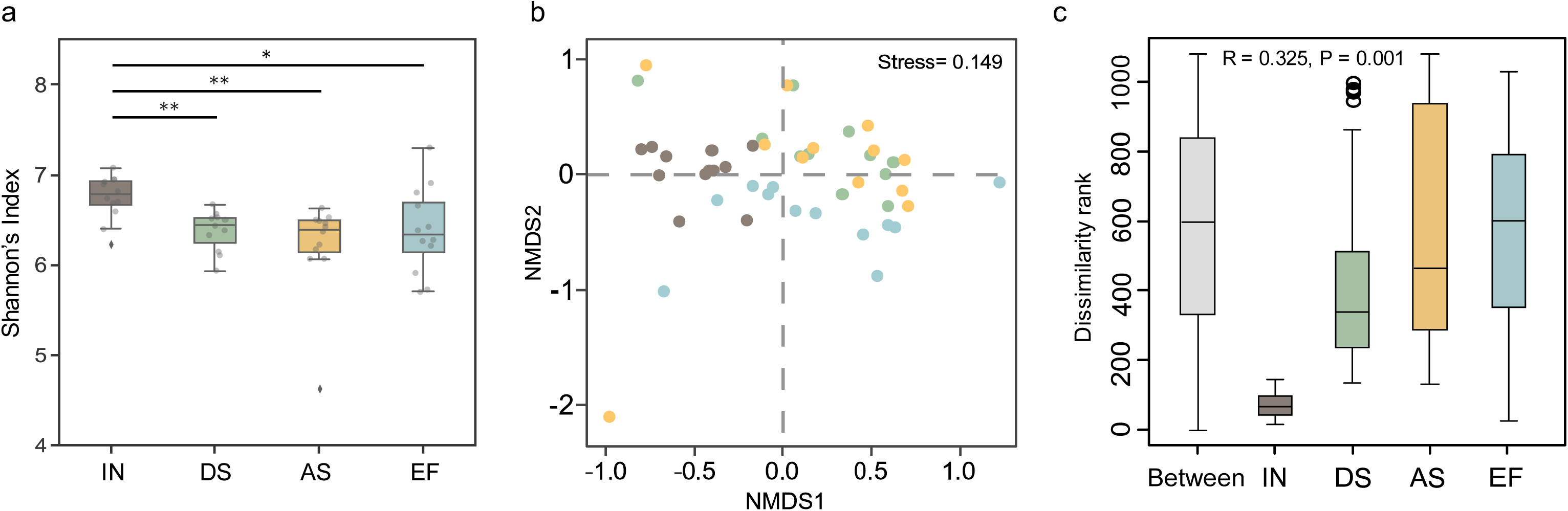
α- and β-diversity of RNA vOTUs throughout the wastewater treatment process. a. α-diversity of RNA vOTUs measured in Shannon’s index across 4 sample types in 47 WWTP samples from Switzerland. b. NMDS analysis of the abundance distribution of RNA vOTUs in 47 WWTP samples from Switzerland. c. ANOSIM analysis showing the dissimilarity rank of RNA vOTU compositions between and within different WWTP compartments in Switzerland.

**Supplementary Fig. 2.**
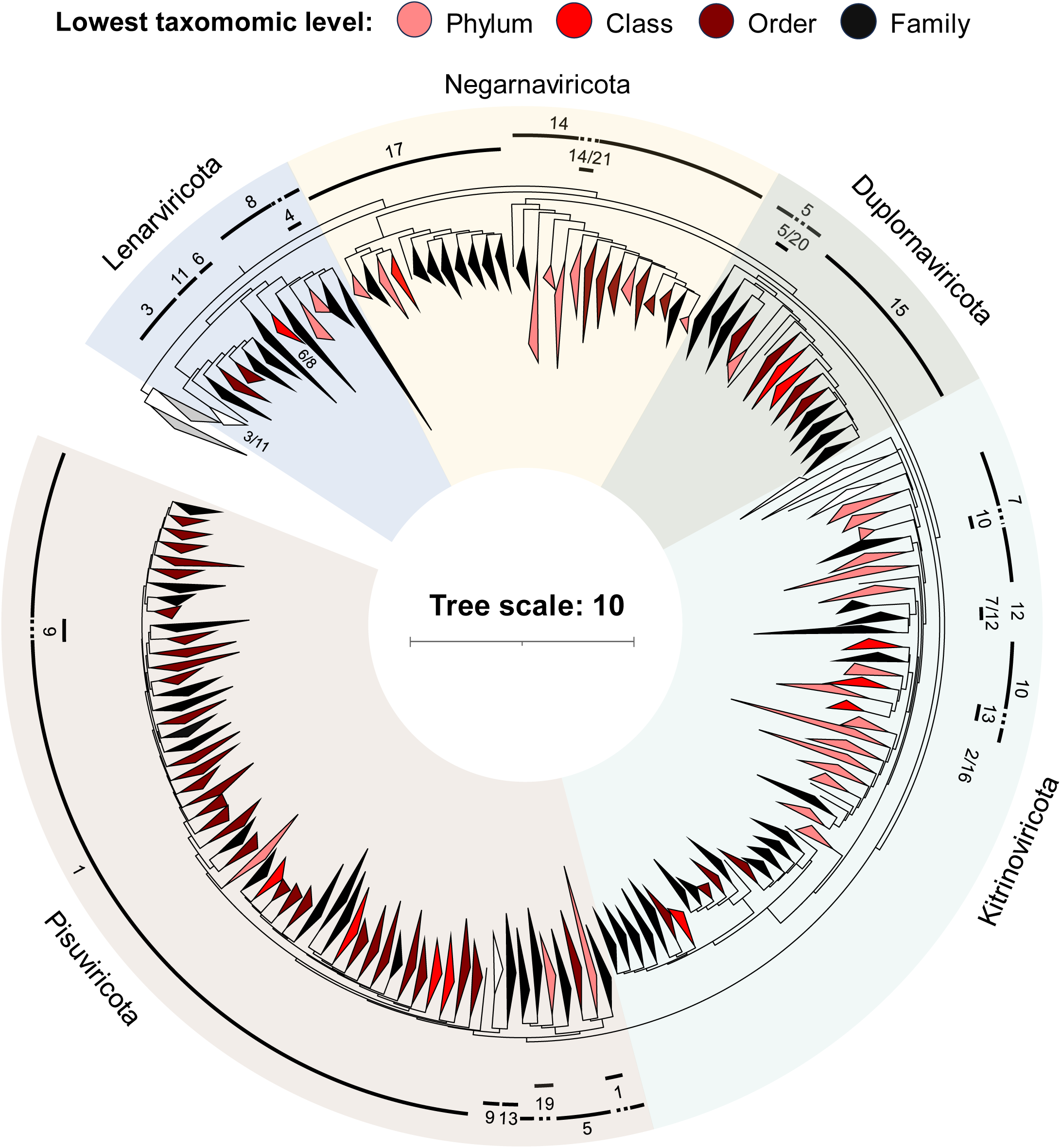
Complete global RdRp phylogenetic tree. The phylogenetic tree includes of all reference RdRp centroids and RNA vOTUs encoding “complete” RdRp domains. The color of each clade in the tree represented the lowest taxonomic level assigned to that clade. The number next to each clade indicates that the majority of its members belong to the corresponding megataxon. Specific labels for each megataxon are provided in Supplementary Table 4.

**Supplementary Fig. 3.**
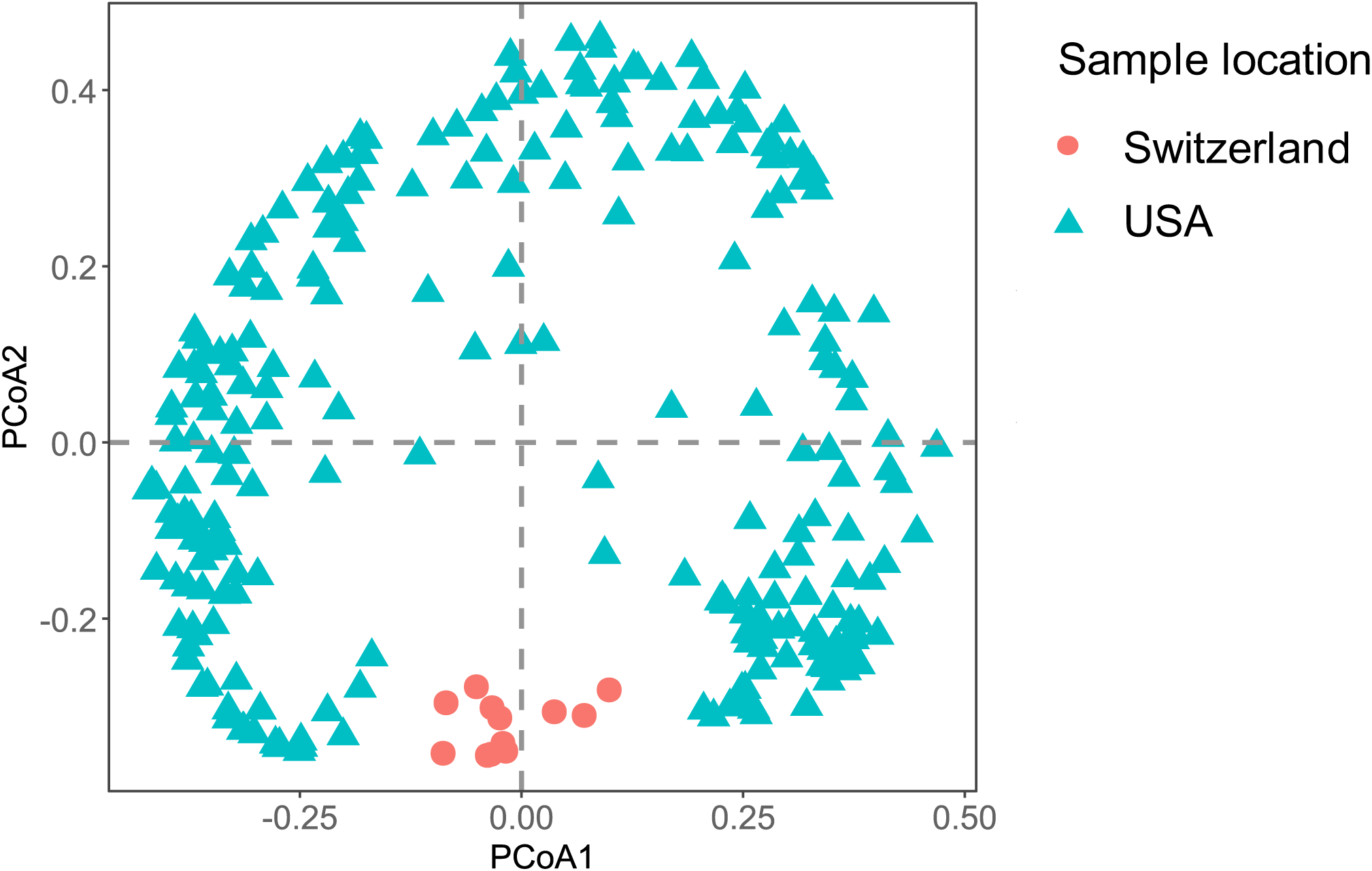
NMDS analysis of the short reads-based distribution of human RNA virus in influent samples from Switzerland and the USA.

**Supplementary Fig. 4.**
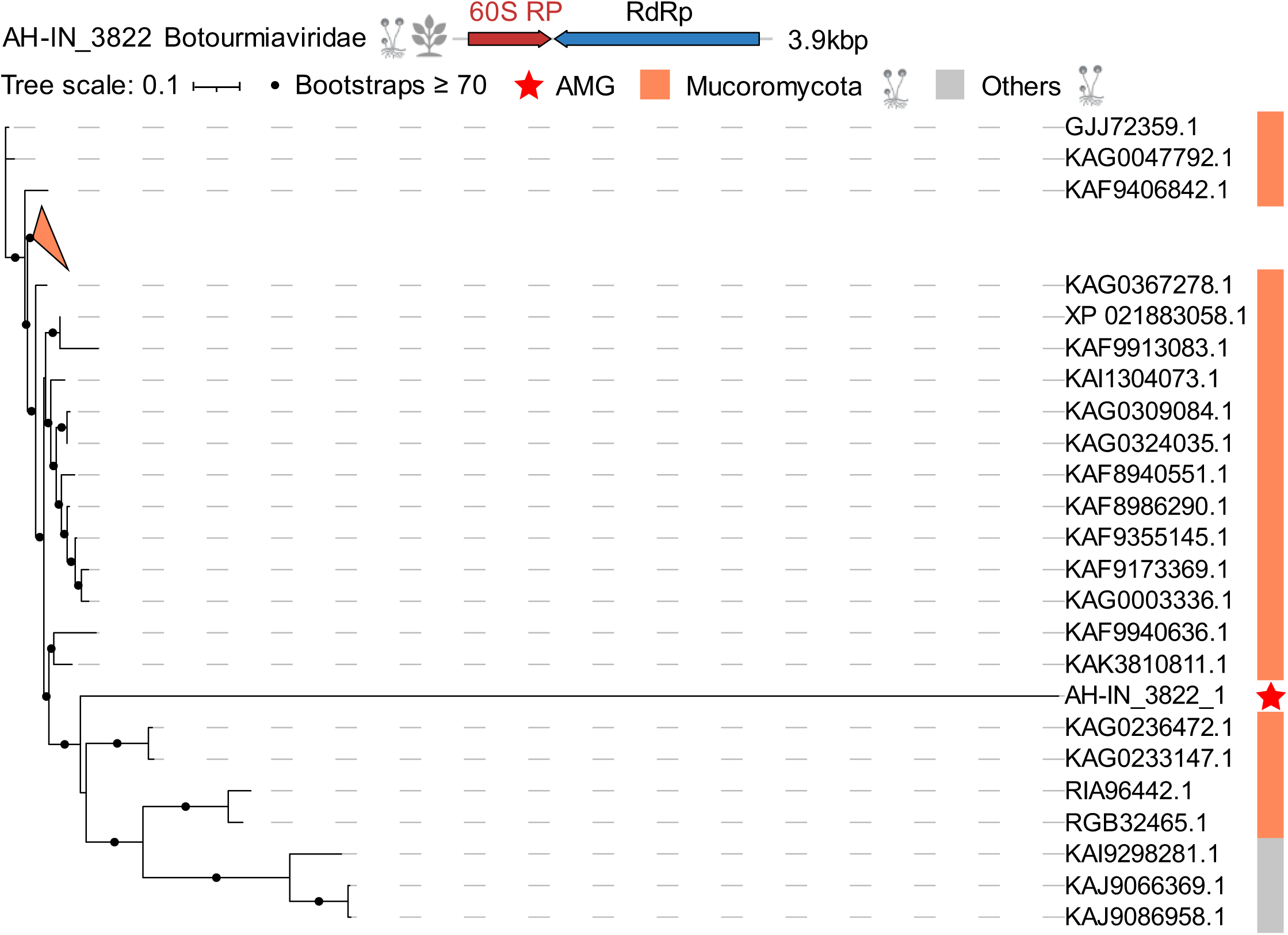
Phylogenetic tree of RNA virus-encoded 60S ribosomal protein and reference sequences from the NCBI nr database.

**Supplementary Fig. 5.**
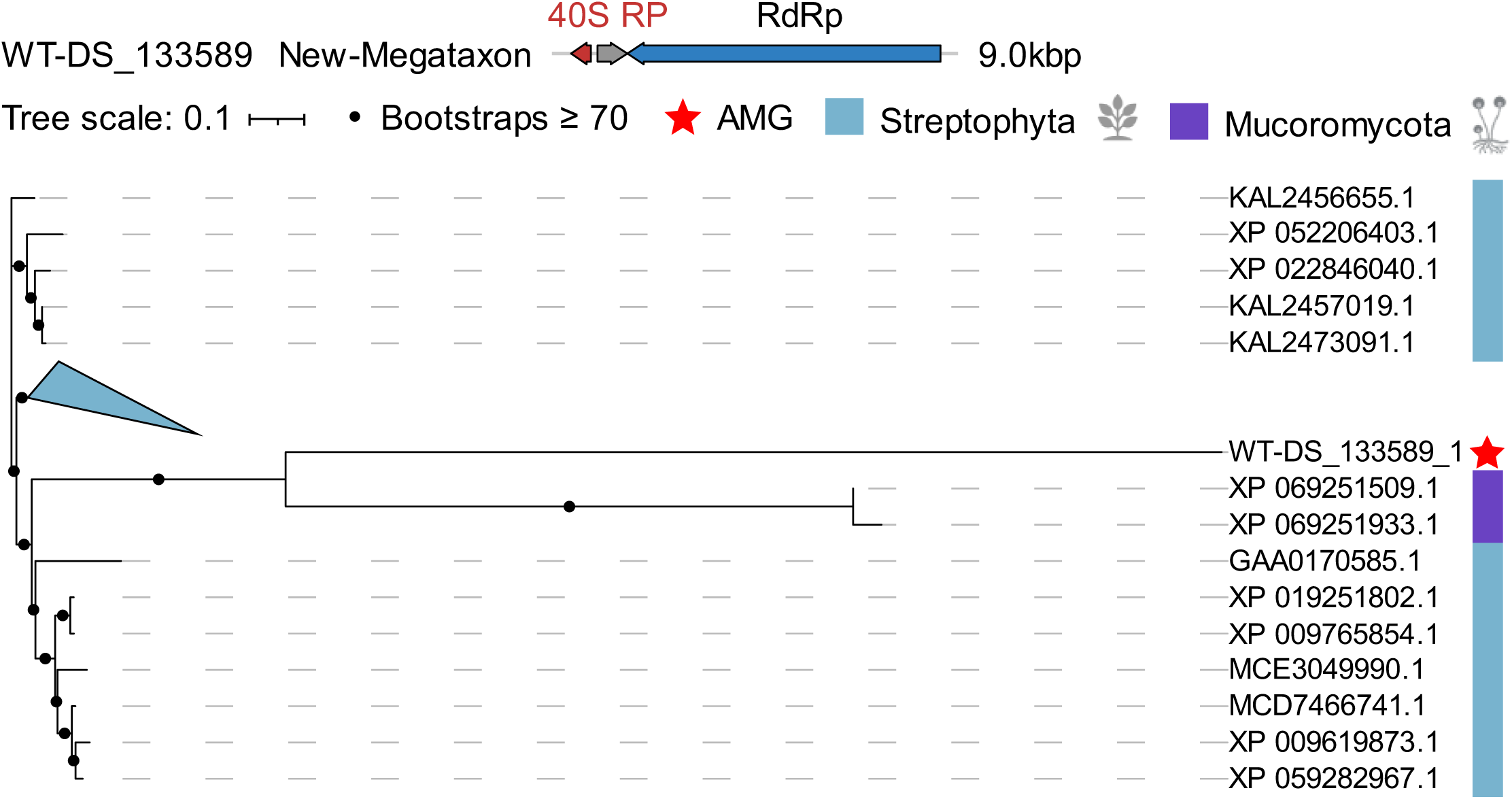
Phylogenetic tree of RNA virus-encoded 40S ribosomal protein and reference sequences from the NCBI nr database.

**Supplementary Fig. 6.**
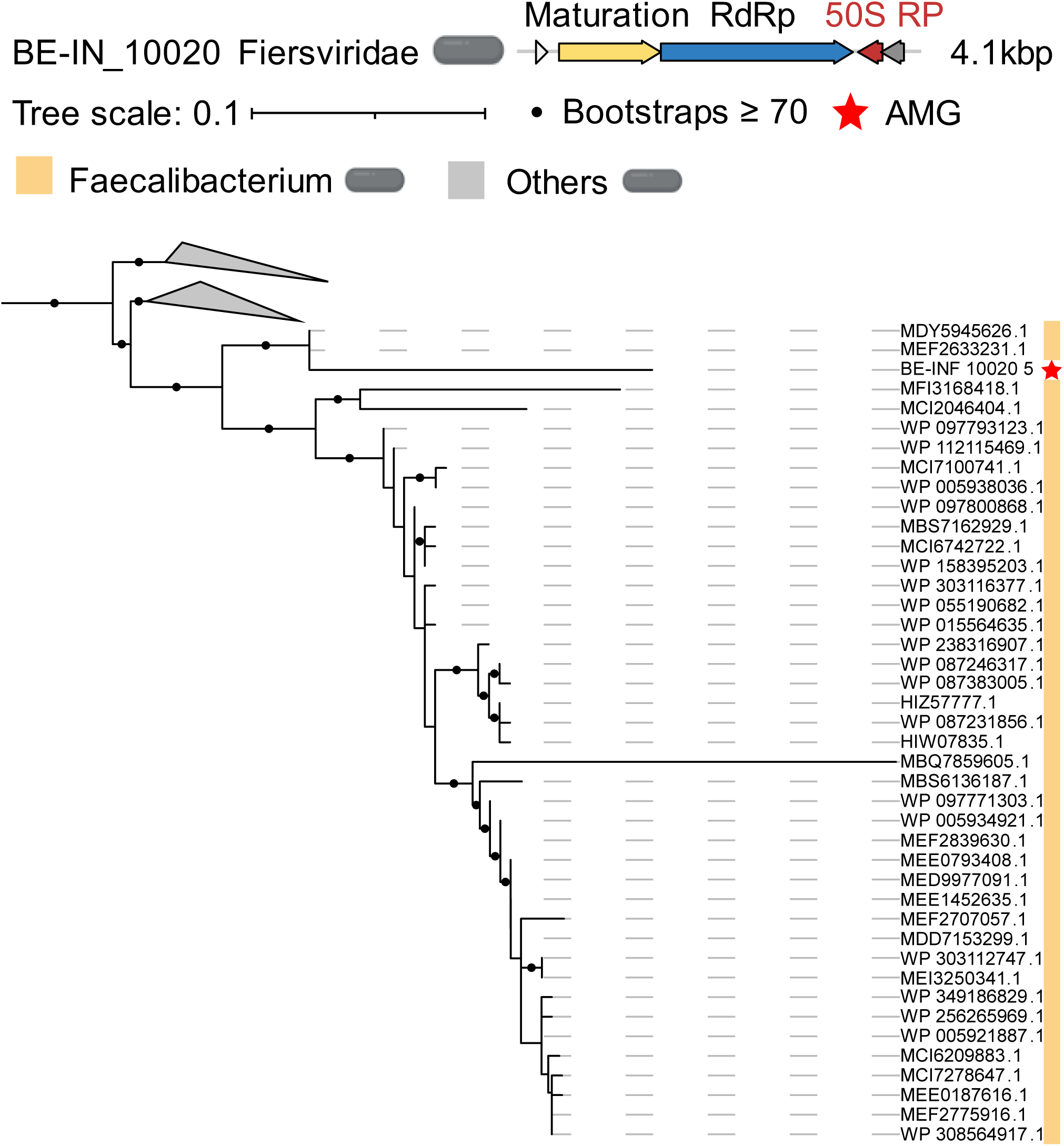
Phylogenetic tree of RNA virus-encoded 50S ribosomal protein and reference sequences from the NCBI nr database.

**Supplementary Fig. 7.**
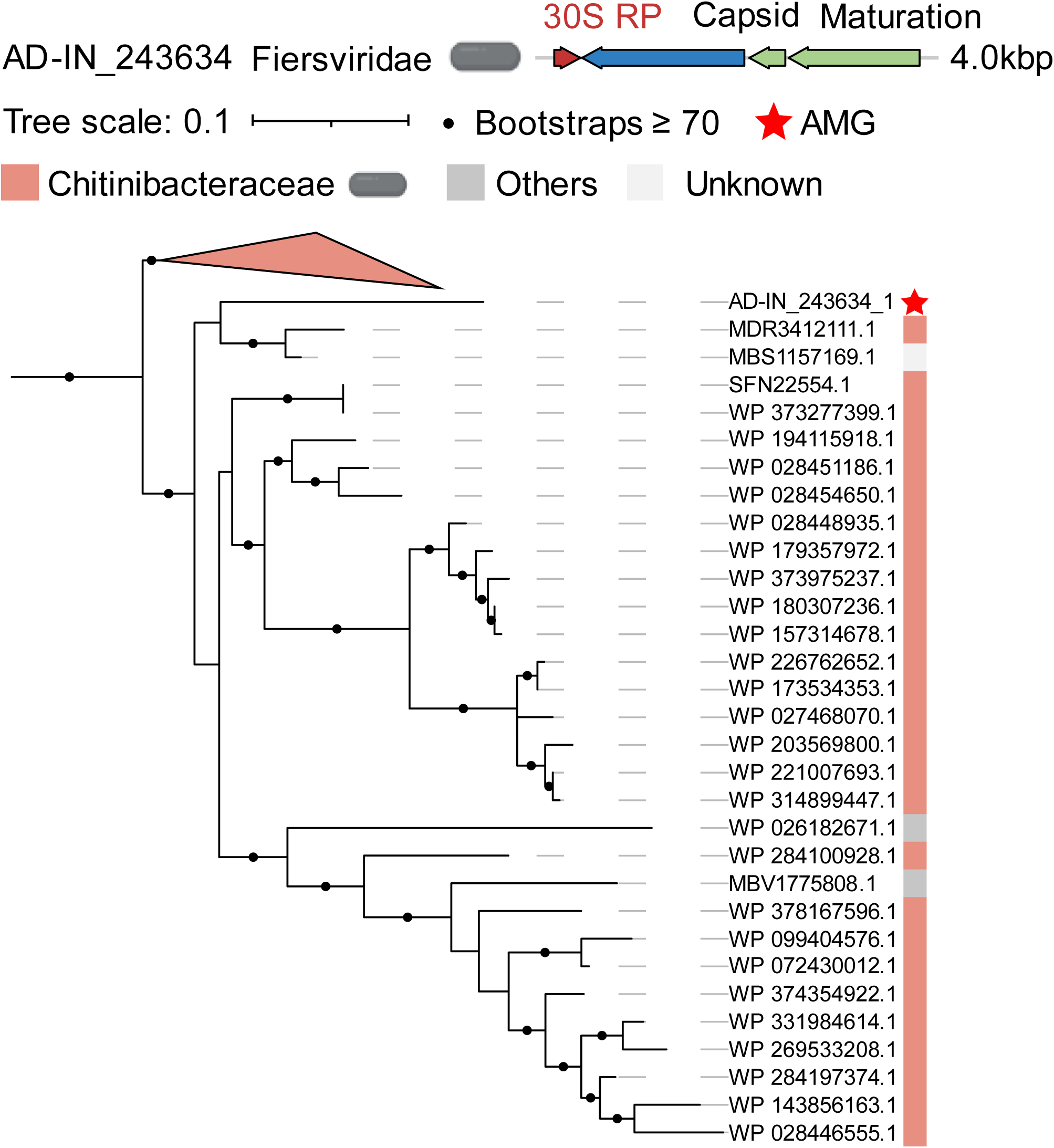
Phylogenetic tree of RNA virus-encoded 50S ribosomal protein and reference sequences from the NCBI nr database.

**Supplementary Fig. 8.**
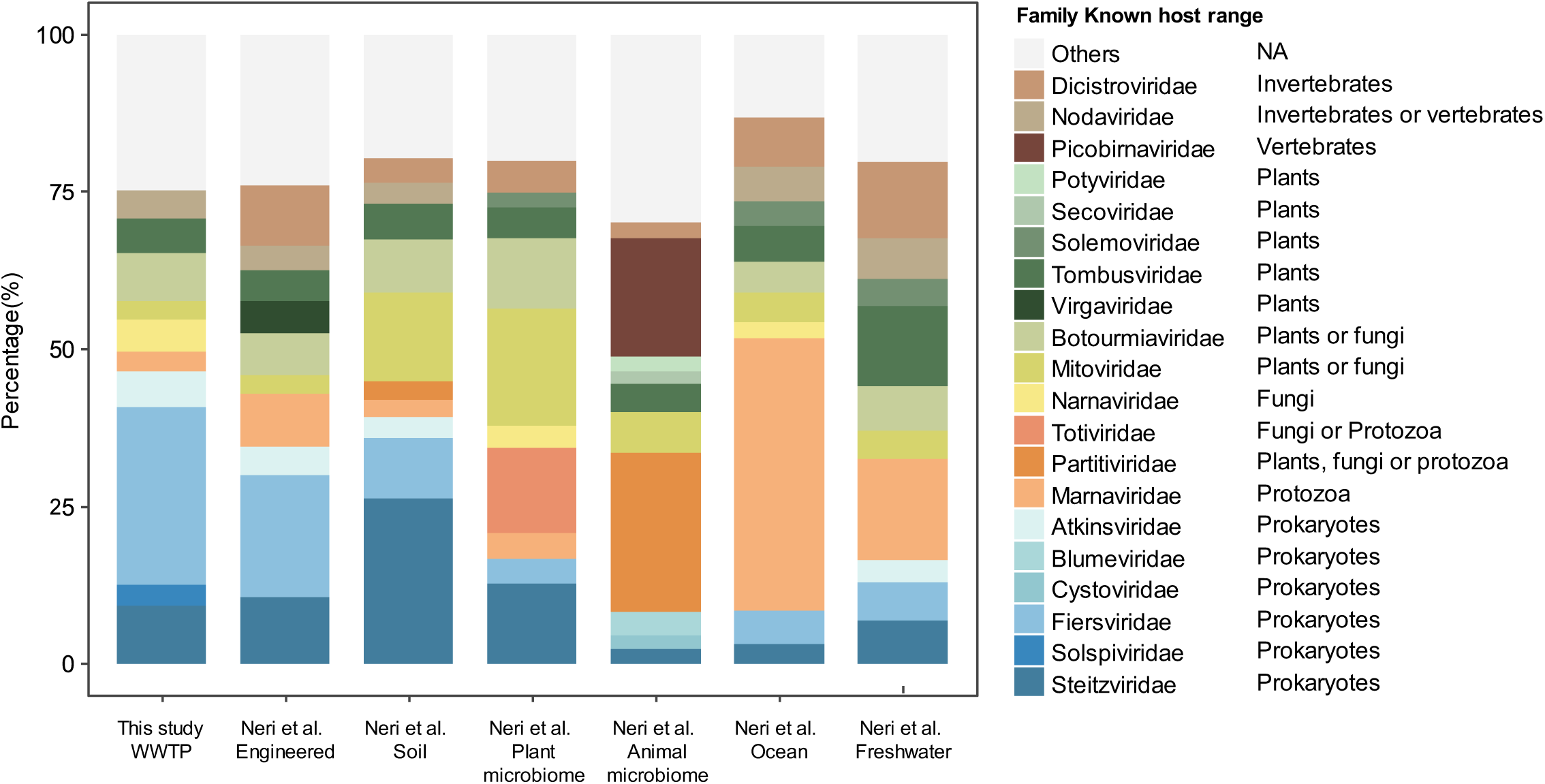
Abundance proportions of the top ten most abundant RNA viral families in this study and other ecosystems surveyed by Neri et al. The proportion of each viral family is calculated by dividing the number of sequences from that family by the total number of virus sequences with family-level classification recovered from the ecosystem.

## References

1. Wu, D., Wu, T., Liu, Q. & Yang, Z. The SARS-CoV-2 outbreak: What we know. International Journal of Infectious Diseases 94, 44–48 (2020).

2. Nimgaonkar, I., Ding, Q., Schwartz, R.E. & Ploss, A. Hepatitis E virus: advances and challenges. Nature Reviews Gastroenterology & Hepatology 15, 96–110 (2017).

3. Hall, A.J. et al. Norovirus Disease in the United States. Emerging Infectious Diseases 19, 1198–1205 (2013).

4. Moreno-Altamirano, M.M.B., Kolstoe, S.E. & Sánchez-García, F.J. Virus Control of Cell Metabolism for Replication and Evasion of Host Immune Responses. Frontiers in Cellular and Infection Microbiology 9 (2019).

5. Gualdoni, G.A. et al. Rhinovirus induces an anabolic reprogramming in host cell metabolism essential for viral replication. Proceedings of the National Academy of Sciences 115 (2018).

6. Dominguez-Huerta G, Zayed A A, Wainaina J M & al., e. Diversity and ecological footprint of Global Ocean RNA viruses. Science 376, 1202–1208 (2022).

7. Shi, M. et al. The evolutionary history of vertebrate RNA viruses. Nature 556, 197–202 (2018).

8. Shi, M. et al. Redefining the invertebrate RNA virosphere. Nature 540, 539–543 (2016).

9. Kamitani, M., Nagano, A.J., Honjo, M.N., Kudoh, H. & Kümmerli, R. RNA-Seq reveals virus–virus and virus–plant interactions in nature. FEMS Microbiology Ecology 92 (2016).

10. Zayed A A, Wainaina J M, al., e. & Sullivan, M.B. Cryptic and abundant marine viruses at the evolutionary origins of Earth’s RNA virome. Science 376, 156–162 (2022).

11. Wolf, Y.I. et al. Doubling of the known set of RNA viruses by metagenomic analysis of an aquatic virome. Nat Microbiol 5, 1262–1270 (2020).

12. Neri, U. et al. Expansion of the global RNA virome reveals diverse clades of bacteriophages. Cell 185, 4023–4037 e4018 (2022).

13. Chen, Y.-M. et al. RNA viromes from terrestrial sites across China expand environmental viral diversity. Nature Microbiology 7, 1312–1323 (2022).

14. Callanan, J. et al. Expansion of known ssRNA phage genomes: From tens to over a thousand. Science Advance 6, eaay5981 (2020).

15. Hou, X. et al. Using artificial intelligence to document the hidden RNA virosphere. Cell 187, 6929–6942.e6916 (2024).

16. Urayama, S.I. et al. Double-stranded RNA sequencing reveals distinct riboviruses associated with thermoacidophilic bacteria from hot springs in Japan. Nat Microbiol 9, 514–523 (2024).

17. Zhang, Y.-Z., Chen, Y.-M., Wang, W., Qin, X.-C. & Holmes, E.C. Expanding the RNA Virosphere by Unbiased Metagenomics. Annual Review of Virology 6, 119–139 (2019).

18. Starr, E.P., Nuccio, E.E., Pett-Ridge, J., Banfield, J.F. & Firestone, M.K. Metatranscriptomic reconstruction reveals RNA viruses with the potential to shape carbon cycling in soil. Proc Natl Acad Sci U S A 116, 25900–25908 (2019).

19. Ju, F. & Zhang, T. Bacterial assembly and temporal dynamics in activated sludge of a full-scale municipal wastewater treatment plant. ISME J 9, 683–695 (2015).

20. Wu, L. et al. Global diversity and biogeography of bacterial communities in wastewater treatment plants. Nat Microbiol 4, 1183–1195 (2019).

21. Ju, F., Xia, Y., Guo, F., Wang, Z. & Zhang, T. Taxonomic relatedness shapes bacterial assembly in activated sludge of globally distributed wastewater treatment plants. Environ Microbiol 16, 2421–2432 (2014).

22. Yuan, L. & Ju, F. Potential Auxiliary Metabolic Capabilities and Activities Reveal Biochemical Impacts of Viruses in Municipal Wastewater Treatment Plants. Environmental Science & Technology 57, 5485–5498 (2023).

23. Shi, L.D. et al. A mixed blessing of viruses in wastewater treatment plants. Water Res 215, 118237 (2022).

24. Guajardo-Leiva, S., Chnaiderman, J., Gaggero, A. & Díez, B. Metagenomic Insights into the Sewage RNA Virosphere of a Large City. Viruses 12 (2020).

25. Karthikeyan, S. et al. Wastewater sequencing reveals early cryptic SARS-CoV-2 variant transmission. Nature 609, 101–108 (2022).

26. Yousif, M. et al. SARS-CoV-2 genomic surveillance in wastewater as a model for monitoring evolution of endemic viruses. Nature Communications 14 (2023).

27. Roux, S. et al. Ecogenomics and potential biogeochemical impacts of globally abundant ocean viruses. Nature 537, 689–693 (2016).

28. Chen, L.X. et al. Large freshwater phages with the potential to augment aerobic methane oxidation. Nat Microbiol 5, 1504–1515 (2020).

29. Yuan, L. et al. Seasonal succession, host associations, and biochemical roles of aquatic viruses in a eutrophic lake plagued by cyanobacterial blooms. Environment International 193 (2024).

30. Sun, M., Yuan, S., Xia, R., Ye, M. & Balcázar, J.L. Underexplored viral auxiliary metabolic genes in soil: Diversity and eco-evolutionary significance. Environmental Microbiology 25, 800–810 (2023).

31. Ju, F. et al. Wastewater treatment plant resistomes are shaped by bacterial composition, genetic exchange, and upregulated expression in the effluent microbiomes. ISME J 13, 346–360 (2019).

32. Orf, G.S. et al. Metagenomic Detection of Divergent Insect- and Bat-Associated Viruses in Plasma from Two African Individuals Enrolled in Blood-Borne Surveillance. Viruses 15 (2023).

33. Levy, J.I., Andersen, K.G., Knight, R. & Karthikeyan, S. Wastewater surveillance for public health. Science 379, 26–27 (2023).

34. Bolger, A.M., Lohse, M. & Usadel, B. Trimmomatic: a flexible trimmer for Illumina sequence data. Bioinformatics 30, 2114–2120 (2014).

35. Li, D., Liu, C.M., Luo, R., Sadakane, K. & Lam, T.W. MEGAHIT: an ultra-fast single-node solution for large and complex metagenomics assembly via succinct de Bruijn graph. Bioinformatics 31, 1674–1676 (2015).

36. Hyatt, D. et al. Prodigal: prokaryotic gene recognition and translation initiation site identification. BMC Bioinformatics 11, 119 (2010).

37. Finn, R.D., Clements, J. & Eddy, S.R. HMMER web server: interactive sequence similarity searching. Nucleic Acids Res 39, W29–37 (2011).

38. Tse, H., Skewes-Cox, P., Sharpton, T.J., Pollard, K.S. & DeRisi, J.L. Profile Hidden Markov Models for the Detection of Viruses within Metagenomic Sequence Data. PLoS ONE 9 (2014).

39. Finn, R.D. et al. The Pfam protein families database: towards a more sustainable future. Nucleic Acids Res 44, D279–285 (2016).

40. Langmead, B. & Salzberg, S.L. Fast gapped-read alignment with Bowtie 2. Nat Methods 9, 357–359 (2012).

41. Rice, P., Longden, I. & Bleasby, A. EMBOSS: the European molecular biology open software suite[<emboss.pdf>. Trends in genetics 16, 276–277 (2000).

42. Steinegger, M. & Söding, J. MMseqs2 enables sensitive protein sequence searching for the analysis of massive data sets. Nature Biotechnology 35, 1026–1028 (2017).

43. Edgar, R.C. Search and clustering orders of magnitude faster than BLAST. Bioinformatics 26, 2460–2461 (2010).

44. Enright, A.J., Van, D.S. & Ouzounis, C.A. An efficient algorithm for large-scale detection of protein families. Nucleic Acids Res 30, 1575–1584 (2002).

45. Edgar, R.C. MUSCLE: multiple sequence alignment with high accuracy and high throughput. Nucleic Acids Res 32, 1792–1797 (2004).

46. Capella-Gutiérrez, S., Silla-Martínez, J.M. & Gabaldón, T. trimAl: a tool for automated alignment trimming in large-scale phylogenetic analyses. Bioinformatics 25, 1972–1973 (2009).

47. Price, M.N., Dehal, P.S. & Arkin, A.P. FastTree 2--approximately maximum-likelihood trees for large alignments. PLoS One 5, e9490 (2010).

48. Letunic, I. & Bork, P. Interactive Tree Of Life (iTOL): an online tool for phylogenetic tree display and annotation. Bioinformatics 23, 127–128 (2007).

49. Nguyen, L.-T., Schmidt, H.A., von Haeseler, A. & Minh, B.Q. IQ-TREE: A Fast and Effective Stochastic Algorithm for Estimating Maximum-Likelihood Phylogenies. Molecular Biology and Evolution 32, 268–274 (2015).

50. Makarova, K.S. et al. Evolutionary classification of CRISPR–Cas systems: a burst of class 2 and derived variants. Nature Reviews Microbiology 18, 67–83 (2019).

51. Bland, C. et al. CRISPR recognition tool (CRT): a tool for automatic detection of clustered regularly interspaced palindromic repeats. BMC Bioinformatics 8, 209 (2007).

52. Nurk, S., Meleshko, D., Korobeynikov, A. & Pevzner, P.A. metaSPAdes: a new versatile metagenomic assembler. Genome Res 27, 824–834 (2017).

53. Steinegger, M. et al. HH-suite3 for fast remote homology detection and deep protein annotation. BMC Bioinformatics 20 (2019).

54. Liu, L. et al. High-quality bacterial genomes of a partial-nitritation/anammox system by an iterative hybrid assembly method. Microbiome 8 (2020).

55. Berlin, F. Mason – A Read Simulator for Second Generation Sequencing Data.<mason201009.pdf>. Technical Report (2010).

56. Lin, Z. et al. Evolutionary-scale prediction of atomic-level protein structure with a language model. Science 379, 1123–1130 (2023).

57. van Kempen, M. et al. Foldseek: fast and accurate protein structure search. Biorxiv 2022**.02. 07.479398** (2022).

58. Schrodinger, LLC (2015).

59. Dixon, P. VEGAN, a package of R functions for community ecology. Journal of Vegetation Science 14, 927–930 (2003).

60. Kang, D.D. et al. MetaBAT 2: an adaptive binning algorithm for robust and efficient genome reconstruction from metagenome assemblies. PeerJ 7, e7359 (2019).

61. Parks, D.H., Imelfort, M., Skennerton, C.T., Hugenholtz, P. & Tyson, G.W. CheckM: assessing the quality of microbial genomes recovered from isolates, single cells, and metagenomes. Genome Res 25, 1043–1055 (2015).

62. Chaumeil, P.A., Mussig, A.J., Hugenholtz, P. & Parks, D.H. GTDB-Tk: a toolkit to classify genomes with the Genome Taxonomy Database. Bioinformatics 36, 1925–1927 (2019).

63. Cambuy, D.D., Coutinho, F.H. & Dutilh, B.E. Contig annotation tool CAT robustly classifies assembled metagenomic contigs and long sequences. bioRxiv (2016).

64. Milanese, A. et al. Microbial abundance, activity and population genomic profiling with mOTUs2. Nat Commun 10, 1014 (2019).

65. Chen, L.-X., Anantharaman, K., Shaiber, A., Eren, A.M. & Banfield, J.F. Accurate and complete genomes from metagenomes. Genome Research 30, 315–333 (2020).

