## Supplementary Texts for "Global RNA Virome in Wastewater Treatment Plants Reveals Ecological Insights and Human Health Implications"

**Legends**

**Supplementary Texts**

Supplementary Text 1. Wastewater effluent samples displayed higher diversity variations

Supplementary Text 2. The relative abundance patterns of RNA vOTUs with different host ranges varied across sample types and countries

Supplementary Text 3. Refinement of global RdRp phylogenetic tree through WWTP RNA vOTUs

Supplementary Text 1: Wastewater effluent samples displayed higher diversity variations

To fairly assess the diversity and abundance of RNA vOTUs throughout the WWTPs, samples from four compartments (i.e., IN, DS, AS, and EF) of 12 WWTPs from Switzerland were selected and comparatively analyzed (47 samples in total). The result showed that the RNA vOTUs in influent samples had significantly higher α-diversity (Fig. S1a). The RNA viral community structures differed distinctly among influent, sludge, and effluent samples (Fig. S1b). Except pairwise relationship between DS and AS (ANOSIM P > 0.05), the RNA vOTUs composition in any two compartments of the WWTPs were significantly different (ANOSIM P < 0.01) (Fig. S1c and Supplementary Table 3). The distinct RNA vOTUs distribution across different compartments indicated that the wastewater treatment process may exert environmental filtering impacts on RNA viruses. The RNA viral community structure of influent samples exhibited the highest similarity among compartments (Fig. S1c). Subsequently, dissimilarities within groups increased as the wastewater progressed through the denitrification and activated sludge tanks, culminating in the highest dissimilarity observed within the effluent sample group (Fig. S1c). Additionally, effluent samples displayed larger variations than influent samples in terms of α-diversity and combined absolute abundance of RNA vOTUs (Fig. S1a and Fig. 1e).

Supplementary Text 2: The relative abundance patterns of RNA vOTUs with different host ranges varied across sample types and countries

Putative prokaryotic RNA vOTUs accounted for more than 60% average abundance proportion of RNA viral community in each WWTP compartment group (i.e., IN, DS, AS, EF) in Switzerland (Extended Data Fig. 3). Putative plants or fungi-associated RNA vOTUs dominated 277 influent and 14 effluent samples from the USA (Fig. 3b). Interestingly, in the activated sludge samples (n = 61) from all the sampled countries, the putative prokaryotic RNA vOTUs was the most abundant category in the majority (n = 53) of samples with no geographical restrictions (Fig. 3b). Anaerobic digester sludge samples showed a distinctly higher proportion of RNA vOTUs that putatively infect protozoa compared with other sample types (Fig. 3b).

Supplementary Text 3: Refinement of global RdRp phylogenetic tree through WWTP RNA vOTUs

For the five major branches of RNA viruses, *Lenarviricota* took the basal position, *Pisuviricota* and *Kitrinoviricota* were placed on the top branch (Fig. 2). Although a previous study^12^ suggested that *Flasuviricetes* may be an “offender” and was placed inside *Pisuviricota*, both megataxon clustering and phylogenetic analysis suggested *Flasuviricetes* should be affiliated into *Kitrinoviricota*, which was consistent with other previous studies^10, 11^. *Cystoviridae* was discovered to be a plausible “offender” again^12^, the whole members from the megataxon of *Cystoviridae* were away from their respective phyla *Duplornaviricota* but posited inside *Pisuviricota*. Besides, after phylogenetic analysis, the large number of family-unclassified WWTP RNA vOTUs further emphasizes the need for the establishment of new RNA virus taxonomies in the future.
